# Long-read single-cell genome, transcriptome and open chromatin profiling links genotype to phenotypes

**DOI:** 10.1101/2025.09.08.674950

**Authors:** Alexandra Pančíková, Ruben Cools, Marios Eftychiou, Margo Aertgeerts, Joris Vande Velde, Heidi Segers, Jan Cools, Luuk Harbers, Jonas Demeulemeester

## Abstract

Current single-cell multiomics methods typically provide limited genomic information, constraining genotype-phenotype studies. To address this gap, we developed SPLONGGET (Single-cell Profiling of LONG-read Genome, Epigenome, and Transcriptome), which integrates 10X Genomics barcoding with Oxford Nanopore sequencing to simultaneously profile genome, chromatin accessibility, and full-length transcriptomes in thousands of single cells. By retaining all tagmentation fragments during library preparation, SPLONGGET delivers whole-genome coverage, supports target enrichment for effective single-cell genotyping, and remains backwards compatible with existing short-read workflows. SPLONGGET enables comprehensive calling of small variants, structural variants, and copy number alterations. Applying SPLONGGET to paediatric B-cell acute lymphoblastic leukaemia revealed clonal dynamics and the phenotypic effects of somatic variants. Notably, we evidence parallel evolution of immune escape variants with four distinct splice site mutations and loss of heterozygosity in the CAR T-cell therapy *CD19* target. In conclusion, SPLONGGET enables integrated high-throughput analysis of genetic variation and molecular phenotypes using off-the-shelf kits, offering a timely and powerful tool to study genetically heterogeneous samples, such as tumours but also ageing normal tissues.

## Introduction

Recent years have seen the emergence of several single-cell multiomic approaches^1^. Currently, the most widely used method is the commercial 10X Genomics Single Cell Multiome ATAC + Gene Expression assay, which enables simultaneous 3’ gene expression and open chromatin profiling of thousands of single cells^2^. Most approaches however provide only limited insights into the genome. Those which do are typically lower-throughput plate-based methods, such as G&T-seq^3^, DNTR-seq^4^, SDR-seq^5^ or scONE-seq^6^. Furthermore, it has become clear that both tumours and normal tissues are genetic mosaics, with complex mixtures and dynamics of evolving clones^7^. To study these systems, and the effects of genetic variation on phenotypes and plasticity, high-throughput single-cell multiome data with a comprehensive genomic component are essential^8^. Such data allows to generate detailed genotype-to-phenotype maps and understand tissue evolutionary dynamics in health and disease.

Like single-cell multiomics, long-read sequencing has become more widespread in recent years. However, the vast majority of single-cell multiomic methods still rely on short-read sequencing. While short-read sequencing is highly accurate and cost-effective, it is unable to capture full-length transcripts or the full range of genetic variation, especially in complex genomic regions. Both mainstay long-read technologies, PacBio and Oxford Nanopore Technologies (ONT) sequencing, are making inroads into single-cell gene expression analysis and their unique capabilities fuel method development^9^. For instance, Nanopore sequencing is uniquely independent of fragment-length and effectively samples molecules from the library solution. Taken together, incorporating long read technologies into single-cell workflows will likely yield novel biological insights.

Acute lymphoblastic leukaemia (ALL) is the most common childhood cancer, with B-cell ALL (B-ALL) representing the majority of cases. Among its subtypes, high hyperdiploid B-ALL is the most prevalent in children^10,11^. This subtype is characterized by high levels of aneuploidy, which likely arises in a pre-leukemic B cell during foetal development. At diagnosis, this leukaemia often shows extensive heterogeneity, with different subclonal populations containing distinct mutations in driver genes such as *NRAS/KRAS*, *FLT3* and *NSD2*^11,12^. Over the past decades, treatment of B-ALL has evolved significantly^13^. Notably, the development of anti-CD19 chimeric antigen receptor (CAR) T-cell therapy has marked a major advancement in the treatment of relapsed/refractory B-ALL. Nevertheless, nearly 50% of patients relapse, most within two years after receiving CAR T-cell therapy^14^. CD19-positive relapse generally arises from loss of CAR T-cell persistence or potency, while CD19-negative relapse may result from antigen escape through acquired (epi)genetic alterations or lineage switching to acute myeloid leukaemia^13,15,16^. To better understand how these complex malignancies evolve and develop resistance, technologies are required which provide simultaneous and high-throughput single-cell genomic, epigenomic, and transcriptomic insights.

To address this gap and study the evolutionary dynamics of genetically heterogeneous samples, we developed Single-cell Profiling of LONG-read Genome, Epigenome, and Transcriptome (SPLONGGET). SPLONGGET relies on widely available 10X Genomics droplet-based barcoding and fragment-length independent Nanopore sequencing. We show how SPLONGGET libraries are backwards compatible with standard short reads and simultaneously allow for deep single-cell genotyping via targeted multiplex PCR and sequencing. We apply SPLONGGET to longitudinal samples from a case of high-hyperdiploid B-ALL throughout therapy and reveal the interplay between genetic variation and molecular phenotypes during somatic evolution.

## Results

### SPLONGGET leverages 10X Genomics barcoding, modified library preparation, and fragment-length independent sequencing to unlock comprehensive single-cell multiomics

Tn5 tagmentation of chromatin in nuclei (as in the Assay for Transposase-Accessible Chromatin sequencing; ATAC-seq) generates fragments ranging from ∼35 base pairs (bp) to several kbp in size^17^. Traditionally, library preparation (notably, bead cleanup and PCR steps) and Illumina sequencing results in only short fragments being read out, providing the typical ATAC-seq open chromatin enrichment. We reasoned that, if the full distribution of fragments could be maintained throughout library preparation and sequenced effectively, this would supplement sparse ATAC signal with whole-genome coverage. With this in mind, we sought to build a novel genome-inclusive single-cell multiome assay based on the widely available 10X Genomics Single Cell Multiome ATAC + Gene Expression assay, which simultaneously tags Tn5-derived fragments and poly(A) mRNA with consistent barcode pairs. We modify the bead cleanups to retain fragments of all sizes, adjust PCR steps to long-range PCR, and omit cDNA fragmentation (**Fig. 1a** and **Methods**). We optimized library preparation on RPMI-8402 cells, ensuring that we capture full length transcripts as well as short and long DNA fragments. As in the standard 10X Genomics Multiome protocol, separate barcoded cDNA and DNA libraries are obtained. In contrast to standard libraries, these display a wide range of fragment sizes: 300 bp to 3500 bp for the cDNA and 200 bp to >10 kb for the DNA (**Supplementary Fig. 1a-b**). To effectively read out these SPLONGGET libraries, we leverage ONT Nanopore sequencing, which is independent of fragment length (**Methods**).

**Figure 1:**
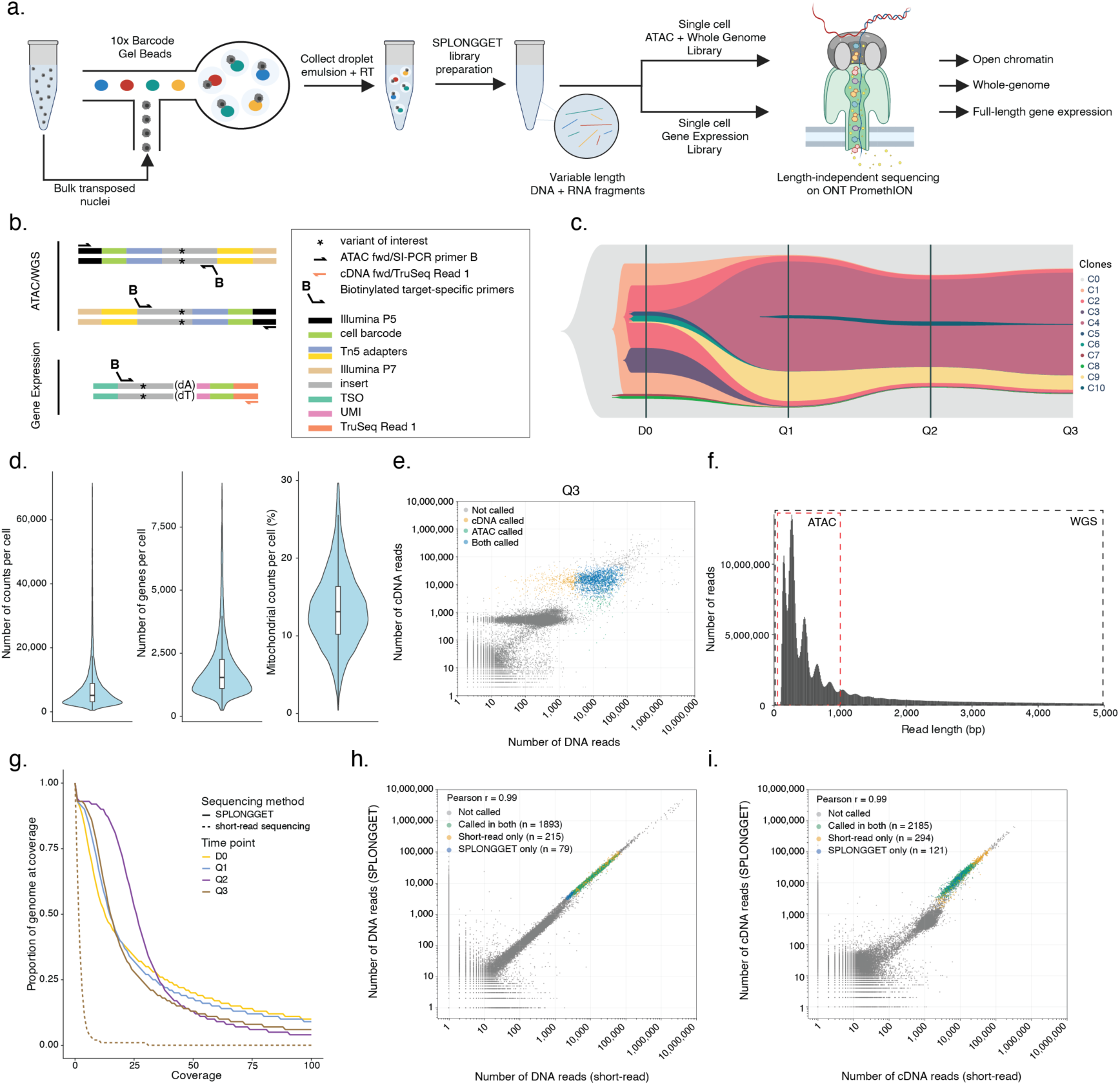
SPLONGGET allows simultaneous long-read profiling of transcriptome, genome and chromatin accessibility. **a**) Schematic of SPLONGGET workflow. **b**) Schematic of targeted SPLONGGET workflow with read structure. **c**) Müller plot depicting clonal composition and evolution of high hyperdiploid B-ALL in XG111 over time. Each coloured area represents a distinct clone, with its relative abundance shown across different time points. C0 denotes the most recent common ancestor (MRCA) from which all subsequent clones are derived. Variants present in other subclones can be found in **Supplementary Table 1**. **d**) Violin plots of high-quality cells after quality control on the cDNA for all time points combined. From left to right: number of UMI counts, genes, and percentage of mitochondrial reads per cell. Boxplot boxes indicate the 25th to the 75th percentile, horizontal bars represent the median, and whiskers extend from –1.5 × IQR to +1.5 × IQR from the closest quartile. IQR, inter-quartile range. **e**) Scatter plot of unique DNA and cDNA read counts for Q3. Colours depict whether the cell passes quality control for ATAC, RNA, or both. **f**) Distribution of read lengths in the genome library. For ATAC analysis, only fragments ≤ 1kbp are used. The right tail of the distribution is cropped for visualization purposes, see **Supplementary Fig. 2f** for the full distribution. **g)** Cumulative coverage distribution for SPLONGGET (D0, Q1, Q2, Q3) and short-read libraries (Q3 only). The y-axis indicates the proportion of the genome covered at or above a given coverage depth (x-axis). **h-i**) Scatter plot of total **h**) DNA and **i**) cDNA reads from Q3 short read and SPLONGGET libraries. Colours depict whether the cell was called in the short reads only, SPLONGGET only, or in both. Pearson correlation was calculated for all barcodes with ≥ 1 count in both libraries.

In addition, we devised an approach to sequence SPLONGGET libraries to saturation in regions of interest. Targeted SPLONGGET aims to provide accurate genotyping of variants of interest, which is crucial for subclonal reconstruction and phylogenetic inference. We reasoned that by taking a fraction of the original untargeted SPLONGGET libraries, performing a multiplex PCR with biotinylated primers flanking regions of interest, and pulling down the desired amplicons using streptavidin beads, DNA and cDNA libraries can be enriched effectively (**Fig. 1b** and **Methods**).

To demonstrate the capabilities of SPLONGGET, we applied it to a case of paediatric B-ALL with a high hyperdiploid karyotype. Multiple subclonal mutations were previously identified in this case through targeted single-cell DNA sequencing with MissionBio Tapestri (patient XG111; **Supplementary Fig. 1c-j**)^11,18^. We applied SPLONGGET to four fresh-frozen bone marrow samples, taken longitudinally at diagnosis (D0), first (Q1), second (Q2), and third relapse (Q3). The patient received standard-of-care chemotherapy. Bortezomib was administered after first relapse, and anti-CD19 CAR T-cell therapy (tisagenlecleucel, Kymriah) was given after second relapse (**Fig. 1c** and **Supplementary Table 1**). After CAR T cell therapy, the patient relapsed again with loss of CD19 protein expression and presence of CAR T cells (CD19-negative B-ALL relapse).

For each time point, we targeted 3,000 cells to balance cell numbers against sequencing costs and ran the cDNA and DNA libraries on two PromethION flow cells each (**Methods**). We obtained on average 175.7 million reads per DNA library and 169.2 million per cDNA library, with an average read length N50 of 2,694 bp and 854 bp for DNA and cDNA, respectively (**Supplementary Fig. 2a-f**). We then processed transcriptomic data using the existing nf-core scnanoseq^19,20^ pipeline and the DNA data using our in-house developed scdnalong workflow^21^. We observed an average duplication rate of 21.1% for the DNA libraries and 41.1% for the cDNA libraries, confirming SPLONGGET library diversity (**Supplementary Fig. 2g, h**). Per standard single-cell best practices, for both library types, cells are called automatically by finding the inflection point from a knee-plot.

The cDNA libraries were further filtered based on total and mitochondrial read counts per cell. Finally, potential doublets were excluded based on total read counts. Cells with >20,000 cDNA UMIs or >60,000 reads in the DNA library were classified as likely doublets and removed from downstream analysis. This was confirmed by assessing single-cell allele counts in a region of clonal loss of heterozygosity (LOH) in the tumour (see below; **Methods**). The resulting 10,106 high-quality cells (D0: 1,808; Q1: 3,897; Q2: 2,051; Q3: 2,350; **Fig. 1d, e**; **Supplementary Fig. 2i-k**, and **Supplementary Fig. 3a-d**) were subsequently used for feature selection, dimensionality reduction, clustering, and annotation (**Methods**).

As expected, genome libraries maintained a wide range of fragment sizes and showed nucleosomal ∼150 bp periodicity (**Fig. 1f and Supplementary Fig. 2f**). For downstream ATAC analysis, reads shorter than 1 kb were used to generate fragment files. After peak calling, cells were further filtered on Transcription Start Site (TSS) enrichment and unique fragments, resulting in 10,268 high-quality cells (D0: 1,995; Q1: 4,942; Q2: 1,327; Q3: 2,004) for chromatin accessibility analysis (**Supplementary Fig. 4a-h** and **Methods**).

We also sought to benchmark SPLONGGET library construction and demonstrate backwards compatibility with standard short-read multiome sequencing. To this end, we fragmented and size selected the SPLONGGET Q3 cDNA library and size selected the SPLONGGET Q3 genome library, as per the standard 10X Genomics Multiome protocol, and performed Illumina sequencing to the recommended depths (**Supplementary Fig. 5a, b** and **Methods**). We ran preprocessing using cellranger-arc followed by the same computational downstream processing and cell type assignment as for long-read libraries. As expected, genome coverage is significantly lower for the short-read library compared to SPLONGGET libraries (e.g., 79-93% covered at ≥ 5X for SPLONGGET vs 6% for short reads, **Fig. 1g**). For both genome and gene expression, barcode read counts and cell calls were highly correlated with those from the long-read SPLONGGET libraries (Pearson correlation: 0.99 for DNA and cDNA, **Fig. 1h, i**). Short-read libraries exhibited highly similar quality control metrics (**Supplementary Fig. 3d**, **Supplementary Fig. 4g, h**, and **Supplementary Fig. 5c, d**). Furthermore, we see a high concordance in clustering, automated cell type assignment (87% of cells with same cell type), and ATAC peaks in marker genes (**Supplementary Fig. 5e-j**). Taken together, SPLONGGET generates high quality single-cell multiome data while adding valuable long-read transcriptome and whole-genome information and retaining maximal compatibility with existing protocols and lab infrastructure.

### SPLONGGET reveals extensive genomic, transcriptomic and regulatory heterogeneity in paediatric B-ALL

To explore the transcriptomic landscape of paediatric B-ALL case XG111, we first sought to characterize the different cell types in our SPLONGGET data. Cell type annotation using celltypist assigned four large clusters as progenitor B-cells, corresponding to the tumour cells of the four time points. The remaining clusters constituted a repertoire of normal immune cell types including (non)-classical monocytes; NK cells; naive B cells; and helper (CD4^+^) and cytotoxic T cells (CD8^+^).

Interestingly, most naive B-cells derive from D0, while they were nearly undetectable at Q3, as may be expected after depletion by the anti-CD19 CAR T-cell treatment. Vice versa, cytotoxic T cells were specific for Q3, likely representing the active CAR T cells (confirmed below). Shifts in the frequencies were also visible for NK cells, monocytes and naive T cells during disease progression (**Fig. 2a-c, Supplementary Fig. 6a-b**, and **Methods**).

**Figure 2:**
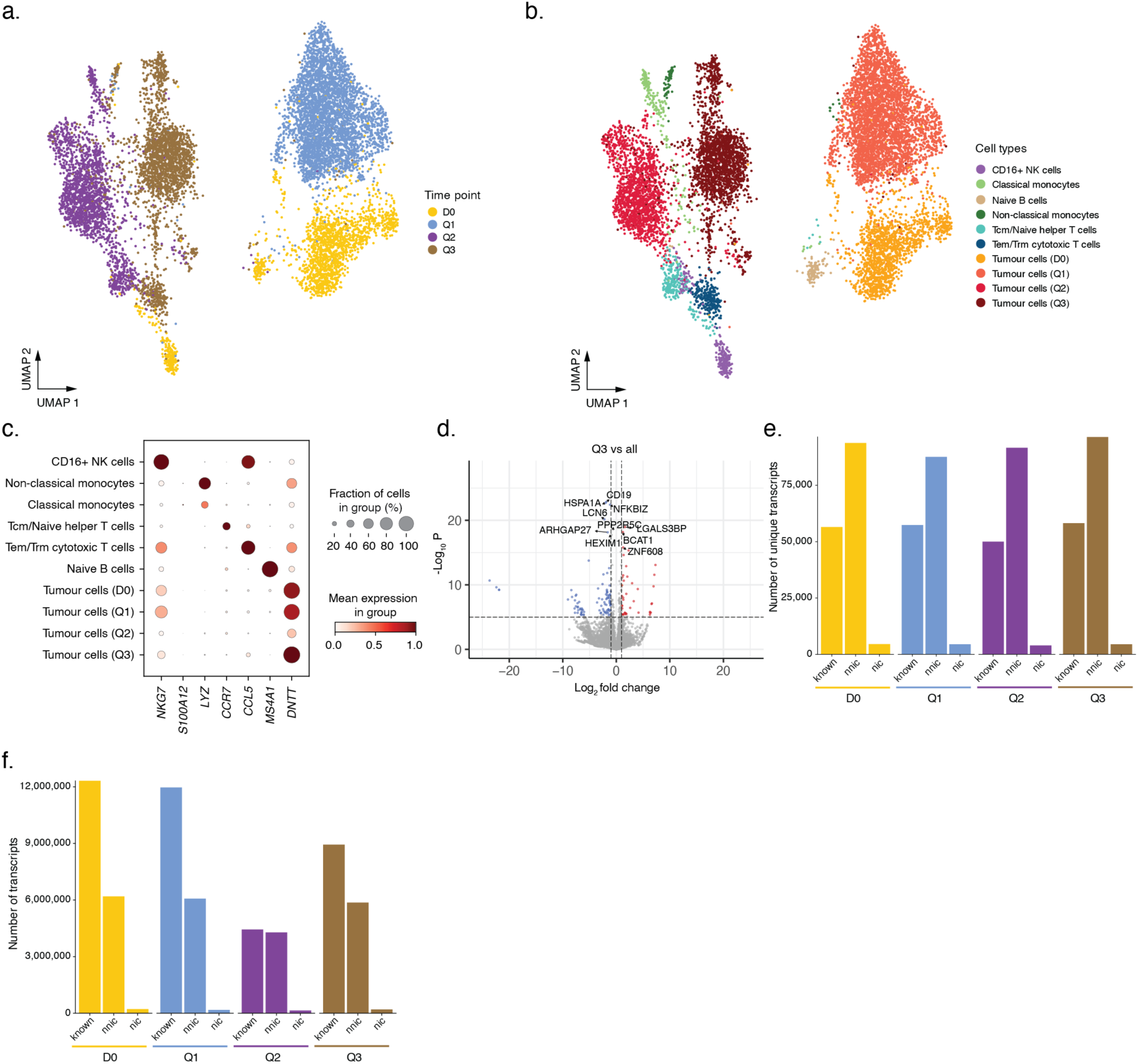
Gene expression landscape of paediatric B-ALL XG111 through therapy and relapse. **a**) UMAP of gene expression data coloured by time point. **b**) UMAP of gene expression data coloured by cell type annotation. **c**) Dot plot of select gene expression markers per cell type. **d**) Volcano plot showing differentially expressed genes at the Q3 time point compared to earlier time points (D0, Q1, and Q2). The top 10 most significant genes are highlighted. **e-f**) Bar plots of **e**) unique and **f**) total transcripts per time point grouped by isoform category; known, nic (novel in catalogue), nnic (novel not in catalogue).

We performed differential gene expression analysis of tumour cells across time points to identify transcriptomic changes within the tumour population. We detected on average 5,530 differentially expressed genes per time point (range: 3,972-7,106; **Supplementary Fig. 6c**). We could confirm downregulation of *CD19* in Q3 relative to the other time points, in line with the clinically observed loss of CD19 protein and CD19-negative relapse (**Fig. 2d** and **Supplementary Fig. 6d-f**). In addition to gene-level analysis, by capturing full-length cDNA, SPLONGGET enables transcript-level analysis and the identification of (novel) isoforms and fusion transcripts. For example, IsoQuant identified 154,867 unique isoforms, of which 98,368 were classified as novel (**Fig. 2e, f**).

To assess gene regulation, we performed pyCisTopic^22^ topic modelling on the SPLONGGET chromatin accessibility profiles (**Fig. 3a**). We leveraged cell type annotations from the transcriptomic layer of our dataset. For cells passing ATAC, but not transcriptome quality controls (2,746 out of 10,268 total cells), we used majority voting between the five nearest neighbours with a transcriptome counterpart (**Methods**). Altogether, we detected 190,760 accessible regions. Clustering based on chromatin accessibility closely recapitulated transcriptome-based clusters (**Fig. 3b**, **Supplementary Fig. 7a**, and **Fig. 2a, b**,). As expected, promoter regions of marker genes showed a similarly cell type-specific increase in chromatin accessibility in the corresponding cluster (**Fig. 3c**). Furthermore, predicted gene activity based on proximal chromatin accessibility supported transcriptome-based cell type annotations (**Fig. 3d**, **Supplementary Fig. 7b-g**, **Methods**). Of note, a small number of Q3 tumour cells, as annotated from the RNA, clusters separately in the ATAC data. The cluster has a uniquely high predicted gene activity of mid-erythroid markers such as *PDRX2*, suggesting a small mid-erythroid population was misclassified based on RNA alone. Its identification highlights a benefit of having transcriptome and chromatin accessibility data from the same single cells.

**Figure 3:**
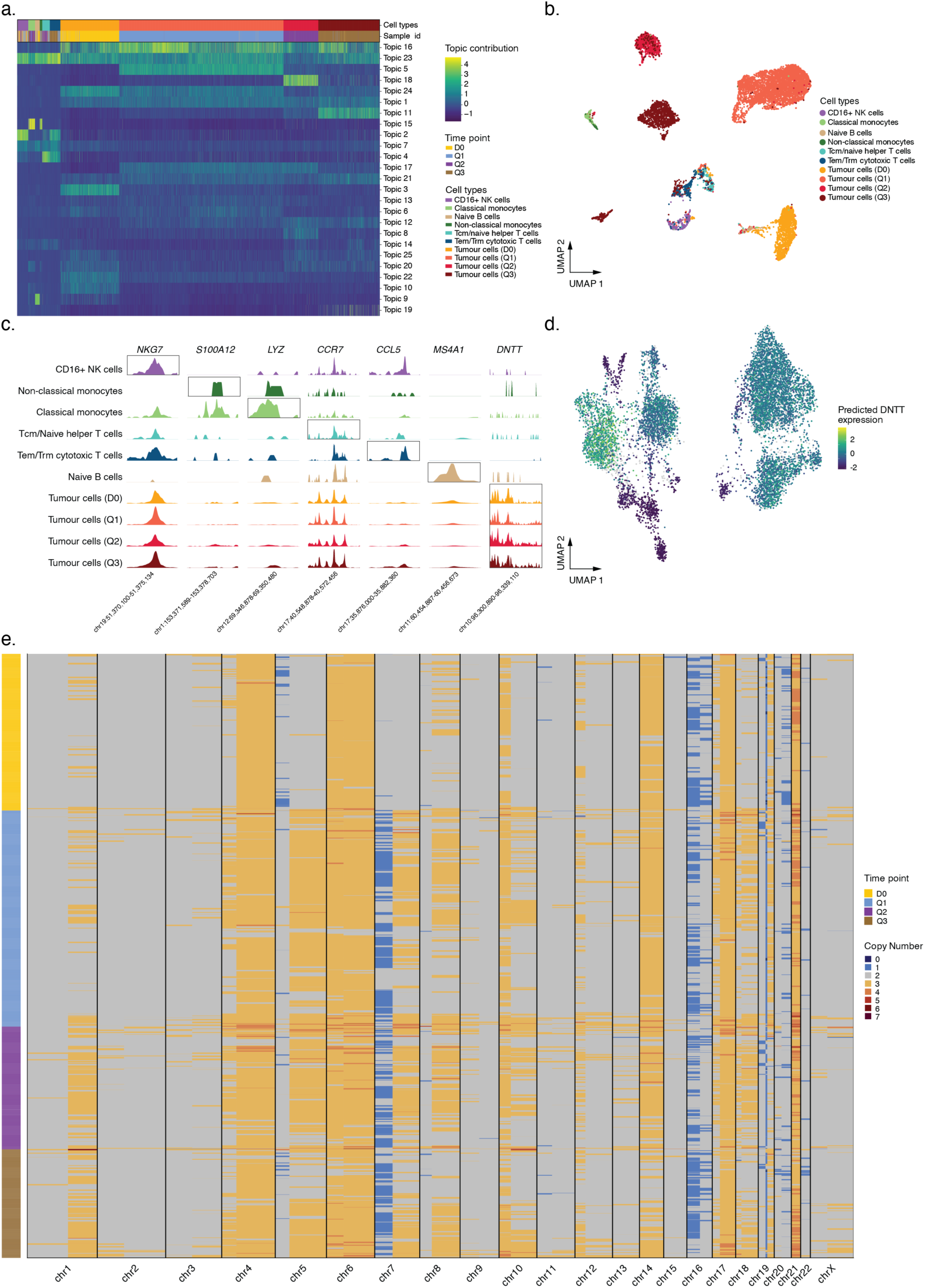
Chromatin accessibility and copy number landscape of XG111. **a**) Heatmap with scaled pyCisTopic cell topic contributions clustered by cell type. **b**) UMAP of chromatin accessibility data coloured by cell type. **c**) Normalised BigWig ATAC pseudobulk profiles for regions surrounding marker genes of different cell types **d**) *DNTT* (tumour marker gene) gene expression prediction based on chromatin accessibility visualized on the gene expression UMAP. **e**) Heatmap of the copy number landscape detected by single-cell DNA-based copy number calling. Each row corresponds to a single cell, with genomic position represented along the x-axis. Total copy number status is colour coded. Cells are hierarchically clustered within each time point.

A key advantage of SPLONGGET is that the genomic library is not limited to highly accessible regions, enabling direct genome-wide detection of copy number alterations (CNAs), structural variants (SVs), single nucleotide variants (SNVs), and indels. First, to investigate CNA heterogeneity and evolution over time, we performed single-cell copy number calling. Even with relatively sparse single-cell coverage (DNA libraries at ∼20% saturation), we derived 3,800 high-quality copy number profiles across the four time points (D0: 1,082; Q1: 1,467; Q2: 596; Q3: 655; **Fig. 3e** and **Supplementary Fig. 7h**). Low-resolution single-cell CNA profiles inferred from the transcriptome data by InferCNVpy, and from previous DNA amplicon-based data^11^, support the accuracy of the SPLONGGET genome-based calls (**Fig. 3e** and **Supplementary Fig. 7i**). As expected, most CNAs were shared between all four time points, including typical gains of chromosome (chr) 6 and 14. Meanwhile, gains of chr 5q and synchronous chr 7p loss and chr 7q gain –suggesting isochromosome formation– only emerged at later time points (**Fig. 3e** and **Supplementary Fig. 7j**). These results demonstrate that SPLONGGET can directly and accurately quantify CNAs without the need to rely on transcriptomic inference. Taken together, single omics layers from SPLONGGET accurately capture a wide array of cell biology, ranging from genomic to regulatory variation and gene expression heterogeneity from the same single cells.

### SPLONGGET enables long-read whole-genome analysis and targeted single-cell genotyping

Understanding the impact of somatic mutations on cell phenotypes is key in understanding how tumours evolve and escape therapy. However, calling variants from single-cell data is challenging, in part due to inherent sparsity. To overcome this, we leveraged our transcriptome-based cell type annotations to create tumour and normal pseudobulks (**Methods**). These can then be used for tumour-normal long-read variant calling, capturing the full spectrum of variants, ranging from SNVs and indels to more complex CNAs and SVs.

First, we generated allele-specific copy number profiles using ASCATv3^23^, extending the single cell-derived CNAs. ASCAT confirms typical features of high hyperdiploid B- ALL cases shared across all time points, including tetrasomy of chr 21, trisomy of chr 4, 6, 10, 14, 17, and 18, and LOH on chr X, in addition to copy neutral LOH of chr 3 (**Fig. 4a, b** and **Supplementary Fig. 8a, b**)^24^. Subsequently, we characterized the SV landscape, identifying 1,362 SVs using Severus (D0: 1,231; Q1: 1,221; Q2: 1,038; Q3: 1,184; **Fig. 4c** and **Methods**). Most SVs were shared among all time points while ∼100 in total were unique to single time points (**Fig. 4e** and **Supplementary Fig. 8c-e**). In parallel, we detect in 28,785 SNVs in total (D0: 14,429; Q1: 16,561; Q2: 17,133; Q3: 18,923) (**Fig. 4d** and **Methods**). Similar to SVs, the majority of SNVs were shared across all four time points, with an increasing number of SNVs observed at the later time points. Together, these data allow a comprehensive view of the genomic landscape of XG111 throughout therapy and relapse (**Fig. 4e** and **Supplementary Fig. 8c-e**).

**Figure 4:**
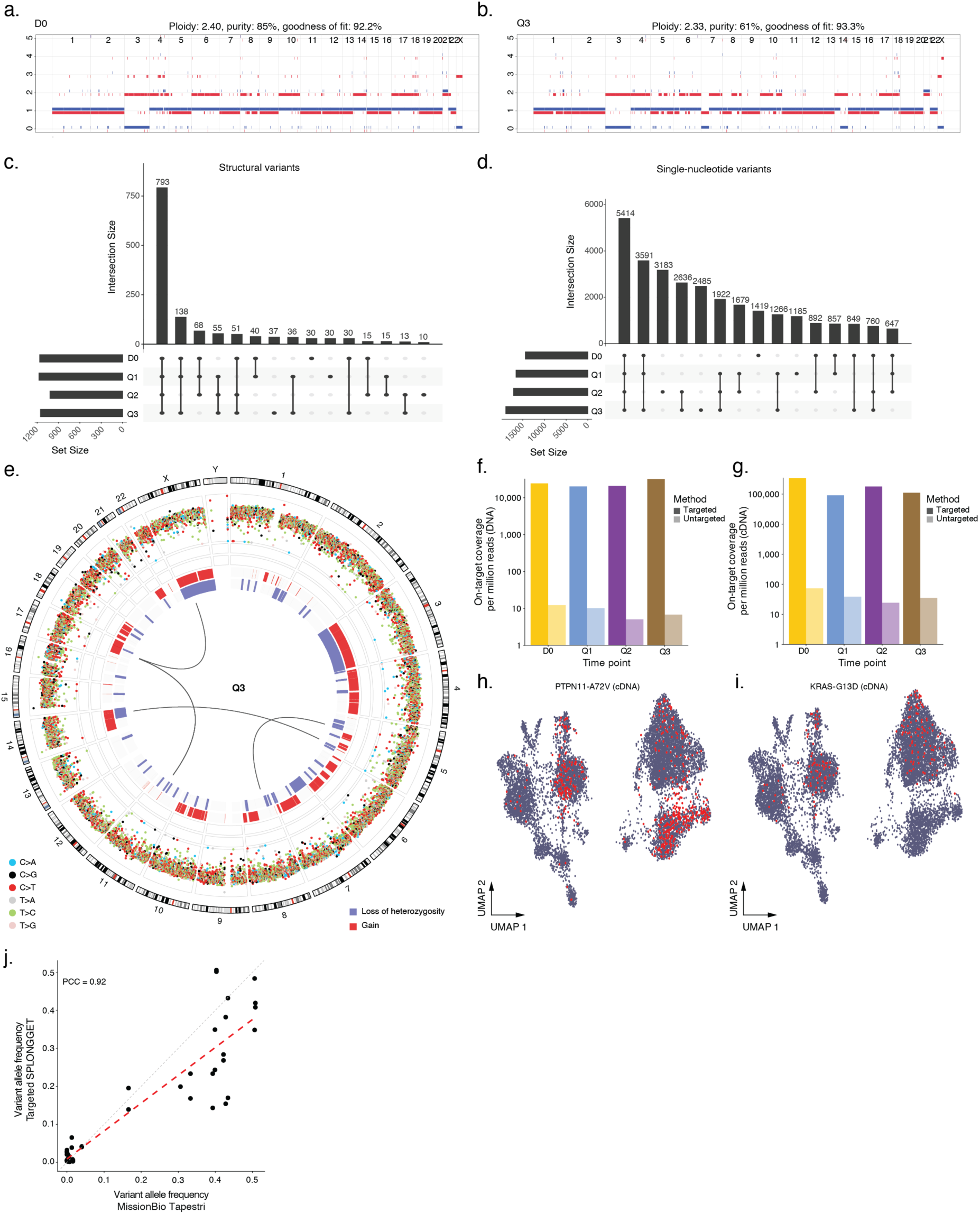
Comprehensive variant calling reveals the mutational landscape of XG111. **a-b**) ASCAT allele-specific copy number profile of XG111 at **a**) diagnosis (D0) and **b**) third relapse (Q3). Red and blue lines indicate the copy number of the major and minor allele, respectively. **c-d**) UpSet plot of **c**) SV and **d**) SNVs detected across the four time points (D0, Q1, Q2, Q3). Each row represents a time point, and each vertical bar corresponds to a specific combination of time points between which the indicated number of variants are shared. **e**) Circos plot of detected genomic alterations at Q3. The innermost track displays translocations. The middle track represents CNAs, where blue bars indicate loss of heterozygosity and red bars indicate gains. The outermost track shows SNVs, colour-coded by substitution class: blue (C > A), black (C > G), red (C > T), grey (T > A), green (T > C), and pink (T > G). **f**-**g**) Coverage of targeted variants for the **f**) DNA libraries and **g**) cDNA libraries. Bars represent on-target coverage per million reads for targeted and untargeted SPLONGGET (shaded). **h-i**) Targeted SPLONGGET (cDNA) of **h**) *PTPN11*^A72V^ and **i**) *KRAS*^G13D^ in single-cells. Cells in which the variant was detected are shown in red on the UMAP. **J**) Dot plot showing the correlation of variant allele frequencies between targeted SPLONGGET (y-axis; mean of cDNA and DNA) and MissionBio Tapestri (x-axis). Each black dot represents a targeted variant in a time point. Dashed red line: linear regression fit. PCC: Pearson’s Correlation Coefficient.

Previous work on XG111 using targeted single-cell DNA sequencing (MissionBio Tapestri) identified (sub)clonal mutations in genes implicated in B-ALL progression (**Supplementary Table 1**, **Figure 1c**)^11^. Specifically, *PTPN11*^A72V^ and *NSD2*^E1099K^ were consistently detected across all time points, while *PTPN11*^E76Q^, *FLT3*^N676K^, *NRAS*^G12D^, and *NRAS*^G13D^ were confined to subclones at D0. *KRAS*^G13D^ was detected mostly at Q1 to Q3 (**Supplementary Table 1**). Mutations present in a sufficient fraction of cells, such as *PTPN11*^A72V^, *NSD2*^E1099K^, and *KRAS*^G13D^ were directly detected during SPLONGGET-based variant calling. (**Supplementary Fig. 8f**). To supplement sparse per cell coverage and boost genotyping of mutations present in only a small subset of cells, we applied target enrichment as described above. This enabled us to sequence our SPLONGGET libraries to saturation in the regions of interest surrounding these variants (**Fig. 1b**, **Supplementary Table 2**). In this targeted approach, we achieve an average 3,242- and 4,402-fold enrichment in coverage at targeted regions for DNA and cDNA libraries respectively (**Fig. 4f, g**). We detect all previously described variants, including rare subclonal ones, and genotype up to 18.4% of cells on average (range 0-75.1%), a 1,272.3-fold increase compared to untargeted SPLONGGET (0.01%, **Fig. 4h, i** and **Supplementary Fig. 9a-j**). Finally, variant allele frequencies observed by targeted SPLONGGET closely match those reported previously (**Fig. 4j** and **Supplementary Fig. 9k**). These results showcase how SPLONGGET enables long-read whole-genome analysis and deep single-cell genotyping at loci of interest through target enrichment.

### SPLONGGET multiomics allows gene regulatory network reconstruction and exploration of genotype–phenotype effects during B-ALL evolution

We used SCENIC+ to integrate SPLONGGET gene expression and open chromatin data and explored gene regulatory network changes in the leukaemic cells over time^22^. We identified direct enhancer-driven regulons (eRegulons) modulated by 29 unique transcription factors (TFs). Several of these TFs exhibited consistently higher expression levels in tumour cells than in normal cells, across all time points (**Fig. 5a**). Notably, dimensionality reduction on the extracted eRegulon data, essentially a biologically informed joint transcriptome-open chromatin embedding, showed separate clusters corresponding to the different cell types (**Fig. 5b**). This confirms that all cell types (or states), including tumour cells from different time points, show specific gene regulatory network activity. For example, *ERG* was more highly expressed in tumour cells compared to normal cells, with its promoter and enhancer regions showing increased accessibility. A member of the erythroblast transformation-specific family and a known oncogene, SCENIC+ predicted that *ERG* co-regulates 1,219 target genes in the leukaemic cells (**Fig. 5c** and **Supplementary Fig. 10a, b**).

**Figure 5:**
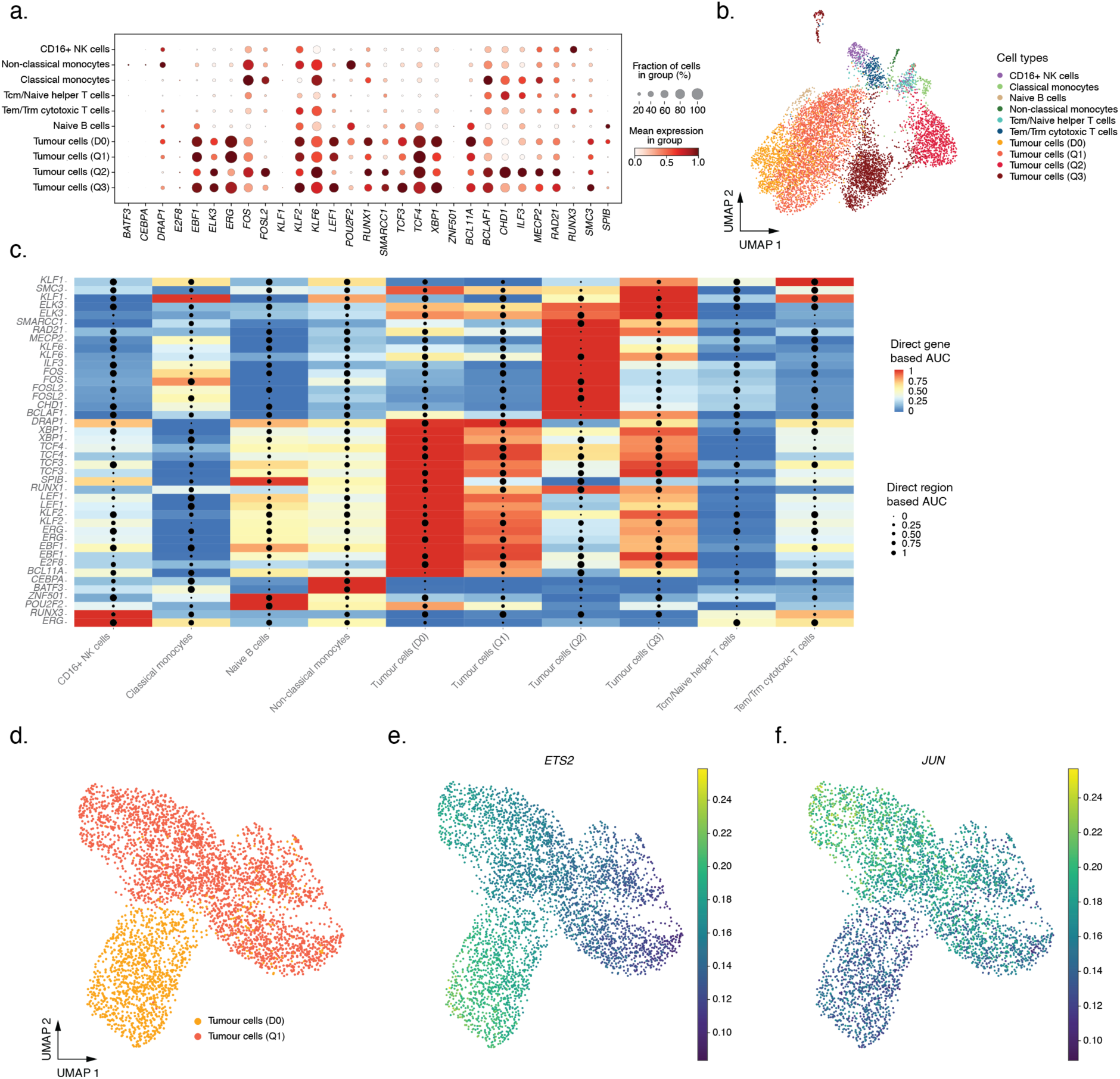
SCENIC+ identifies relevant tumour enhancer–gene regulatory networks. **a**) Dotplot of expression of TFs identified as regulators by SCENIC+. **b**) UMAP projection of eRegulon enrichment scores coloured by cell type. **c**) Heatmap with gene expression and chromatin accessibility enrichment in eRegulons across cell types. **d**) UMAP projection of tumour cells at D0 and Q1 based on eRegulon enrichment scores. **e-f**) UMAP projection with **e**) *ETS2* and **f**) *JUN* eRegulon enrichment score.

Next, we investigated the effect of CNAs on gene expression and chromatin accessibility, integrating the DNA-based single-cell copy number profiles. We focused on chr7, which is diploid at D0 but shows a loss and gain of its p- and q-arms, respectively, in later time points. As may be expected, observed expression and chromatin accessibility are lower on the p-arm in Q1 compared to D0, while they are increased on the q-arm (**Supplementary Fig. 10c, d**). This directly reveals the molecular phenotypic effect of large-scale copy number gains and losses on the measured chromatin accessibility and gene expression. Importantly, inclusion of a simultaneous genomic read-out, allows to study these effects directly and in-depth, instead of relying on circular CNA inference from RNA or open chromatin first. For instance, we hypothesized that the genomic differences between D0 and Q1 tumour cells might be reflected in observed gene regulatory changes. To assess this, we subset 4,178 tumour cells from D0 and Q1 and re-identified eRegulons specifically active in one or the other time point using SCENIC+. We identify the cancer-associated *ETS2* and *JUN* eRegulons, specific to D0 and Q1, respectively (**Fig. 5d-f**). As expected, both eRegulons show a difference in number of observed versus expected target genes on chr5 and 7 (**Supplementary Fig 10e, f**). This demonstrates how the effects of large-scale CNAs can influence inferred gene regulatory network activities. Taken together, integrative analysis of SPLONGGET multiomic data affords uniquely comprehensive insights into the biology of genotypically and phenotypically heterogenous samples.

### Parallel evolution of CAR T-cell escape phenotypes through loss of heterozygosity and splice-site mutations

Finally, to explore the mechanisms of escape from anti-CD19 CAR T-cell therapy, we focused on the Q3 time point. We first sought to identify the CAR construct in our data and performed fusion detection using CTAT-LR on the transcriptome layer. This revealed *TNFRSF9*::*CD247* transcripts restricted to Q3, two genes known to be part of the tisagenlecleucel CAR (CTL019; **Supplementary Table 3**; **Methods**). We wondered if we could also detect the CAR in the genomic data, potentially representing integrated vector copies. Inspection of genomic read coverage in the CAR-constituent gene parts (*CD247/CD3ζ*, *CD8A*, *TNFRSF9*) showed increased coverage in four exons of CD247, with chimeric reads across multiple regions supporting the fusion gene (**Fig. 6a** and **Supplementary Fig. 11a**).

**Figure 6:**
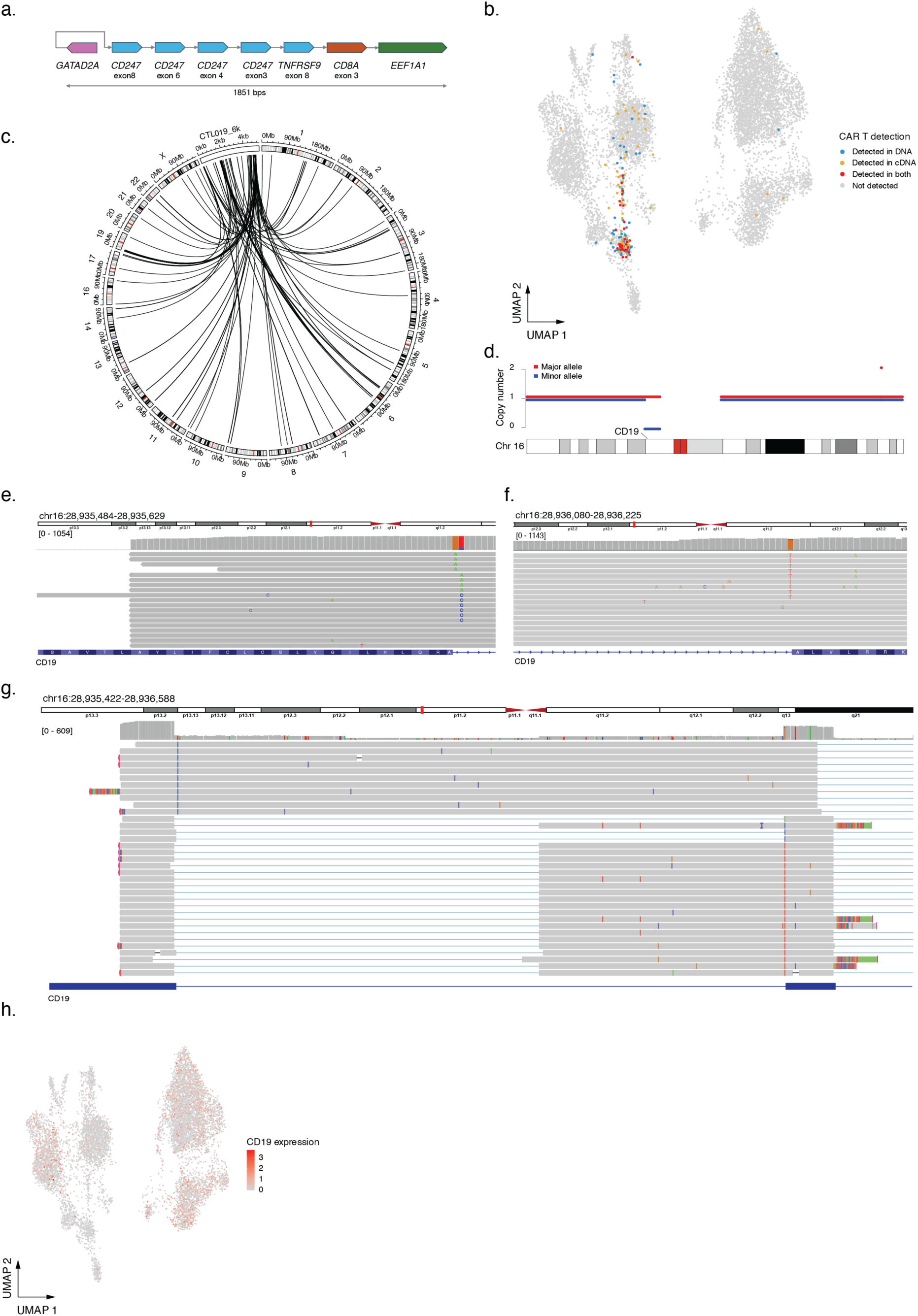
Multiple hits and parallel evolution underpin CD19-negative B-ALL relapse in XG111. **a**) Alignment pictogram visualization of a single long read (read id: CCCGCTAAGATTGAGC_#af6b2432-1b60-445f-891c-221e4e282176_+1of1) derived from the CAR construct and its integration site in an intron of *GATAD2A*. **b**) UMAP showing cells with CAR-derived reads in their genomic data, gene expression data or both. **c**) Circos plot with detected integration sites of the CAR vector (tisagenlecleucel, Kymriah; CTL019). **d**) Zoom in of chr 16 ASCAT allele-specific copy number profile of XG111 at Q3. Red and blue lines indicate the copy number of the major and minor allele, respectively. *CD19* is indicated with a line. **e**) IGV visualisation of Q3 DNA reads at *CD19* exon 4 with the donor splice site with three SNVs. **f**) IGV visualisation of Q3 DNA reads at *CD19* exon 5 with the acceptor splice site with one SNV. **g**) IGV visualisation of Q3 transcripts across exons 4 and 5 of *CD19* evidencing (partial) intron retention as the impact of the splice site mutations. Note the cryptic splice site activation in the intron upon mutation of the original acceptor site. **h**) gene expression UMAP with *CD19* expression.

Motivated by the detection of the CAR construct in both transcriptomic and genomic data we wondered if we could directly genotype cells harbouring this construct and additionally pinpoint vector integration sites. First, we isolated genomic reads reporting part of the CAR and performed *de novo* genome assembly using Flye^25^ to reconstruct the full vector construct as a single contig (**Methods**). We then realigned our genomic and transcriptomic data to the hg38 reference genome including the reassembled CAR contig (CTL019). By identifying reads uniquely mapping to CTL019, we identified all cells in our dataset containing a CAR construct. As expected, CTL019-derived reads were detected in cells from Q3 only and were strongly enriched in the cytotoxic T cell cluster. A large number of cytotoxic T cells had both RNA and DNA reads directly supporting presence and expression of the CAR (**Fig. 6b**). Moreover, supplementary alignments from genomic reads aligned to CTL019 identified vector integration sites across nearly all chromosomes, confirming the polyclonal status of the CAR T-cell population at relapse (**Fig. 6c**).

We next explored the mechanisms underpinning the CD19-negative relapse at Q3. Close inspection of the allele-specific copy number calls revealed a Q3-specific 8Mb deletion on chr16 affecting, among other genes, *CD19* (**Fig. 6d**). In addition to this deletion, we detected three separate SNVs at conserved +1 and +2 positions of the 5’ splice donor site of exon 4, and one more SNV at the -1 position of the 3’ splice acceptor site of exon 5 of *CD19* (chr16:28,935,555 G>A; chr16:28,935,556 T>C; chr16:28,935,556 T>A; chr16:28,936,152 G>T; **Fig. 6e, f**)^26^. Long-read phasing across these sites indicates that all four variants affect distinct copies or subclones. Variant Effect Predictor^27^ confirmed all four variants are likely high impact splice donor or splice acceptor variants (**Supplementary Table 4** and **Methods**). While chr16:28,935,556 T>A was previously reported in a patient with gastrointestinal diffuse large B-cell lymphoma (cosmic ID COSV107329247) and chr16: 28,936,152 G>T was previously reported in patient with common variable immunodeficiency^28^, these variants have not yet been reported in the context of CAR T-cell therapy resistance. The two SNVs with the highest genomic variant allele frequencies were also readily detectable in the transcriptomic data where they are associated with transcripts showing (partial) intron retention (**Fig. 6g**). Specifically, mutation of the 5’ splice site resulted in intron retention while mutation of the 3’ site activated a cryptic splice site in the intron, in both cases likely leading to either nonsense-mediated decay or translation of a truncated *CD19* protein^29^. The combination of LOH and multiple parallel splice site-abrogating SNVs likely underpins the reduced *CD19* expression and protein levels at Q3, resulting in immune escape by several subclones (**Fig. 6h**). These observations suggest parallel evolution of CAR T-cell evasion under strong selection pressures on the B-ALL cells and highlight splice site mutation and LOH as potential contributing factors. Taken together, SPLONGGET long reads support local *de novo* assembly and detection of vector integration sites. Simultaneous detection of variants across multiple ‘omics layers couples genotype and phenotype and, in the case of this B-ALL, uncovers parallel evolution and mechanisms of immune escape.

## Discussion

We have developed SPLONGGET, a single-cell long-read multiome method providing simultaneous genome, open chromatin, and transcriptome data from thousands of single cells. SPLONGGET leverages the widely available and validated 10X Genomics Single Cell Multiome ATAC + Gene Expression kit to tag genomic fragments and transcripts with consistent barcode pairs. Uniquely, by retaining all fragment sizes, as well as full length cDNA, and using ONT Nanopore sequencing, SPLONGGET enables in-depth characterization of single-cell genotypes and molecular phenotypes. On top, we also developed a target-enrichment approach which allows to locally sequence these diverse libraries to saturation and provide deep single-cell genotyping.

Recent years have shown an increase in methodologies to extract transcriptomic and (epi)genomic read outs from the same cell. However, the majority of these methods only profile the transcriptome with either genome or chromatin accessibility, limiting our ability to study the full range of genomic variation and their effects on cellular phenotype^3,4^. Furthermore, current single-cell long-read sequencing methodologies are mostly limited to the transcriptome layer^30^. SPLONGGET was conceived out of a need for high-throughput linking of genotypes-and-phenotypes at the single-cell level. In particular, to study the impact of genetic heterogeneity and somatic evolution in tumours, but also as we have come to appreciate more recently, across normal tissues.

We show that we can reliably detect small variants, allele-specific CNAs, and SVs from the genomic reads. Simultaneous gene expression and open chromatin patterns can effectively guide annotation-based pseudobulking. For instance, in tumour samples with reasonable numbers of both tumour and admixed normal cells, this allows somatic variants to be called in the more accurate tumour-matched normal mode from just a single sample. This avoids additional sampling or cell sorting in cases where a reference normal control is not always taken (e.g., tumour biopsies). Vice versa, tumour cells and tumour-normal doublets can often be unambiguously identified by genotyping identified haplotypes in regions of LOH or other somatic variants.

To demonstrate the rich insights which can be gleaned from SPLONGGET data, we applied it to longitudinal samples from a paediatric case of high hyperdiploid B-ALL through therapy and relapse. We showed that SPLONGGET accurately profiles full-length transcript expression, chromatin accessibility, and genotypes ranging from single-cell CNAs to SVs and small variants. Additionally, we demonstrated that we can locally enrich SPLONGGET libraries to boost genotyping of variants of interest in single cells by performing multiplex PCR with biotinylated primers. Crucially, regions of interest can first be identified by untargeted SPLONGGET followed by target enrichment on the same libraries, as we demonstrate with for *CD19* splice site mutations. By integrating our multiomic data we detected tumour and time point-specific enhancer-gene regulatory networks and the effect of CNAs on these networks. We demonstrate how rich SPLONGGET data can be used creatively to address *ad hoc* questions, for instance genotyping CAR T cells and identifying vector integration sites by *de novo* assembly and read realignment. These findings underscore the enhanced biological insights which can be gained from integrating comprehensive genomic information into a single-cell multiomic approach.

With SPLONGGET, we detailed molecular mechanisms underpinning resistance against anti-CD19 CAR T-cell therapy in high hyperdiploid B-ALL. In XG111, clinically confirmed CD19-negative leukaemic cells likely derived from LOH of *CD19* with multiple subclonal splice site mutations leading to intron retention and unproductive transcripts on the remaining copy. Although *CD19* splice site mutations have been reported in B-ALL relapse after CAR T-cell therapy, the majority of reported resistance mutations are missense and nonsense mutations^15,33,34^. Interestingly, XG111 contains four previously unreported distinct parallel splice site mutations spanning a single intron. These mutations and the additional LOH of *CD19* suggest intense selective pressure on the tumour cells during CAR T-cell therapy. Future studies may shed more light onto whether the extent of parallel evolution and splice site mutation under CAR T-cell therapy seen in XG111 represents an exception or is more widespread.

We further validated SPLONGGET and demonstrated backwards compatibility by using the Q3 library as a starting point for standard short-read multiome profiling (ATAC + gene expression). Both methods result in comparable number of cells being detected, with concordant quality metrics, clustering and cell type annotations. This confirms that SPLONGGET libraries retain all biological information one may expect from a standard short-read 10X Single-Cell Multiome run, and can provide much more if read out with Nanopore long reads. Notably, this offers users flexibility, where they can perform SPLONGGET library preparation and still opt for any combination of short-read or Nanopore long-read sequencing based on their research questions, budgets, or other restrictions.

To run chromatin accessibility computational analysis, we selected DNA reads ≤ 1kbp. However, since all reads theoretically start and end at Tn5 integration sites, one can instead use all tagmentation sites to generate fragment files as demonstrated by Hu et al.^31^. This would allow usage of all the available DNA data for chromatin accessibility profiling. Interestingly, we observed that SPLONGGET chromatin accessibility profiles appear to offer more granular two-dimensional clustering, allowing us to recover cell types or states with few cells, such as the small mid-erythroid population missed in the initial transcriptome annotation.

Given that SPLONGGET requires only a 10X Genomics Chromium device and an ONT Nanopore sequencer, we anticipate it can readily be implemented by a wide array of labs. Similarly, we are convinced it is applicable to a broad range of biological problems including tumour–microenvironment interactions, somatic mosaicism and evolution in normal tissues, but also to study dynamic processes such as development, or isoform-specific gene regulation. While we demonstrated SPLONGGET on fresh frozen human liquid biopsies, we anticipate it can be applied to any sample type that is compatible with the 10X Genomics Multiome approach.

In summary, SPLONGGET is a novel, easily implementable, and backwards compatible method for simultaneous long-read profiling of transcriptome, chromatin accessibility, and whole-genome of thousands of single-cells. Applied to longitudinal samples from a case of paediatric B-ALL, SPLONGGET links genotypes to phenotypes and demonstrates the interplay between ‘omics layers during tumour evolution, through therapy and relapse.

## Materials & Methods

### Sample collection

Bone marrow samples were collected from a case of paediatric HeH B-ALL (XG111) at Leuven University Hospital (UZ Leuven). The study was approved by the Ethics Committee Research UZ / KU Leuven (S68798), with written informed consent obtained in accordance with the Declaration of Helsinki. The patient was treated according to the European Organisation for Research and Treatment of Cancer (EORTC) 58081 protocol. Mononuclear cells were extracted using Ficoll-Paque and were viably frozen. Targeted amplicon-based (MissionBio Tapestri) single-cell sequencing data of XG111 has been published and described before^11^.

### SPLONGGET multiome library preparation

A detailed, step-by-step, protocol is available at protocols.io (https://www.protocols.io/private/697F7B787C1311F0AF690A58A9FEAC02). Briefly, high-quality nuclei isolation from the viably frozen mononuclear cells was performed in accordance with the 10x Genomics Nuclei Isolation for Single Cell Multiome ATAC + Gene Expression Sequencing protocol (CG000365). Following the 10X Genomics Single Cell Multiome Protocol (CG000338), a total of 9,000 nuclei per sample are targeted and processed for transposition. Transposed nuclei were loaded into a single well of a Chromium Next GEM Chip J and processed using the Chromium X instrument to generate Gel Beads-in-emulsion (GEMs). GEMs were collected and incubated to initiate reverse transcription of mRNA and barcoding of transposed DNA fragments. A portion of the resulting emulsion corresponding to a minimum of 3,000 nuclei is broken and used for further custom processing of barcoded cDNA and genomic fragments. The 10X Genomics pre-amplification step was modified to contain: 50 µl LongAmp® Hot Start Taq 2X Master Mix (New England Biolabs, M0533), 4 µl Pre-Amp Primers (10x, PN 2000271), and 46 µl sample. PCR cycling conditions were optimized for LongAmp® polymerase: 65°C for 5 min 95°C for 3 min; 7 cycles of 95°C for 30 sec, 63°C for 6 min; and a final extension at 65°C for 5 min. Post-PCR, a 1.6X SPRIselect bead cleanup (Beckman Coulter, B23318) was performed, eluting in 115 µL Buffer EB (Qiagen, 19086), to retain fragments of all sizes.

Next, the pre-amplified sample, containing barcoded cDNA and genome fragments, was divided into 40 µL for ATAC library construction and 35 uL for cDNA amplification. For ATAC Library Construction, the following reaction was set up: 50 µL LongAmp® Hot Start Taq 2X Master Mix, 4 µL SI-PCR Primer B (10X Genomics, PN-2000128), 2.5 µL individual Sample Index N, Set A (10X Genomics, PN-3000427), and 46 µL pre-amplified sample. PCR cycling was optimized for LongAmp® polymerase: 95°C for 1 min 30 sec; 8 cycles of 95°C for 30 sec, 65°C for 6 min; and a final extension at 65°C for 5 min. Post-PCR, a 1.6X SPRIselect bead cleanup was performed, eluting in 45 µL Buffer EB. For cDNA Amplification, the following conditions were used: 50 µL LongAmp® Hot Start Taq 2X Master Mix, 15 µL cDNA amplification primers (10X Genomics, PN 2000089), 35 µl pre-amplified sample. PCR cycling conditions were as follows: 95°C for 3 min; 7 cycles of 95°C for 20 sec, 63°C for 1 min 30 sec; and a final extension at 65°C for 1 min. Post-PCR, a 0.6X SPRIselect bead size selection and cleanup was performed eluting in 45 uL Buffer EB.

### SPLONGGET target enrichment

Target enrichment starts from standard SPLONGGET single-cell barcoded, non-fragmented cDNA and genome libraries. Custom biotinylated primers (20–25-mers) were designed within 3-300 bp of the target regions (either genomic or transcript coordinates; see Fig. 1b for a schematic overview; **Supplementary Table 2**) with up to five targets multiplexed per PCR. For single-cell barcoded non-fragmented cDNA, 20 ng was used as input with a fixed primer, “Pre-AMP cDNA Fwd,” targeting the TruSeq Read 1 sequence. For the genome libraries, 100 ng was used as input and two reactions were set up, each with one fixed primer, “Pre-AMP ATAC Fwd,” targeting the Illumina P5 sequence and either the forward or reverse primer targeting each region of interest.

First, a pre-pull-down biotin-tagging PCR was performed in 25 µL reactions for both cDNA and genome fragments. Each reaction contained 12.5 µL LongAmp® Hot Start Taq 2X Master Mix, 0.2 µM of each biotinylated primer (up to five per reaction), 0.2 µM Pre-AMP primer (“Pre-AMP cDNA Fwd” or “Pre-AMP ATAC Fwd”), and either 20 ng single-cell barcoded non-fragmented cDNA or 100 ng genome library. PCR cycling conditions for genome fragments were: 95°C for 3 min; 20 cycles of 95°C for 20 sec, a temperature ramp from 57°C to 53°C at −0.4°C/sec for 10 sec, 52°C for 10 sec, 65°C for 4 min; and a final extension at 65°C for 5 min. For non-fragmented cDNA, cycling conditions were: 95°C for 3 min; 20 cycles of 95°C for 20 sec, 57°C for 10 sec, 54°C for 20 s, 65°C for 2 min; and a final extension at 65°C for 5 min. A 1X AMPure XP bead cleanup (Beckman Coulter, A63881) was performed, eluting in 10 µL nuclease-free water.

<u>For Biotin pull-down,</u> 2X and 1X Wash/Bind buffers (5 mM Tris-HCl pH 7.5, 1 M NaCl, 0.5 mM EDTA, nuclease-free water) were prepared. Per PCR reaction, 5 µl (50 µg) M280 Streptavidin Dynabeads (Invitrogen, 11205D) were washed twice with 1 mL 1X Wash/Bind buffer and resuspended in 10 µL 2X Wash/Bind buffer. 10 µL biotinylated cDNA or genome PCR products were mixed with 10 µl (50 µg) washed M280 Streptavidin Dynabeads and incubated on a Hula mixer for 20 min at room temperature. The streptavidin-biotin amplicon conjugate was washed a total of three times with 1 mL 1X Wash/Bind buffer, followed by a final wash with 200 µL 10 mM Tris-HCl pH 7.5. The streptavidin-biotin amplicon conjugate was resuspended in 20 µL nuclease-free water.

To elute fragments for sequencing, a post-pull-down PCR was performed. Each reaction contained 25 µL LongAmp® Hot Start Taq 2X Master Mix, 0.2 µM (up to five) target-specific primer(s), 0.2 µM Pre-AMP primer (“Pre-AMP cDNA Fwd” or “Pre-AMP ATAC Fwd”), 19 µL streptavidin-biotin amplicon conjugate, topped up with nuclease-free water to 50 µL total. PCR cycling conditions mirrored those of the pre-pull-down PCR: for the genome libraries, 95°C for 3 min; 20 cycles of 95°C for 20 s, 57°C to 53°C at −0.4°C/s for 10 s, 52°C for 10 s, 65°C for 4 min; and 65°C for 5 min; for the cDNA, 95°C for 3 min; 20 cycles of 95°C for 20 s, 57°C for 10 s, 54°C for 20 s, 65°C for 2 min; and 65°C for 5 min. Post-PCR, a 1X AMPure XP bead cleanup was performed, eluting in 30 µL Buffer EB

### Nanopore sequencing of (targeted) SPLONGGET libraries

Separate Nanopore sequencing libraries for DNA and non-fragmented cDNA were prepared using the Ligation Sequencing Kit V14 (Oxford Nanopore Technologies, SQK-LSK114). According to the Ligation Sequencing Amplicons V14 protocol, input amounts were adjusted to 150 fmol. Short Fragment Buffer (Oxford Nanopore Technologies, SFB) was used during library preparation wash steps to retain all fragment sizes. For each library, 25 fmol was loaded onto PromethION R10.4.1 flow cells and sequenced for 72 h on a PromethION 24. With a yield of ∼80-100M reads per flow cell, we typically achieve a coverage of ∼4000 transcripts per cell for a library containing ∼3,000 cells. For the DNA libraries, we typically sequence on 2 PromethION flow cells to achieve a mean xxx reads per cell.

### Short read library preparation and sequencing

To validate the SPLONGGET single-cell long-read cDNA and ATAC data, we leveraged the Q3 SPLONGGET libraries and adapted them into short-read Illumina cDNA and ATAC libraries. For the cDNA, 25% of the total amplified, non-fragmented SPLONGGET cDNA library was used as input to fragmentation. This was performed using the 10x Genomics Fragmentation Mix at 32°C for 5 min, followed by end repair and A-tailing at 65°C for 30 min, as specified in the 10X Single Cell Multiome protocol (CG000338). A double-sided SPRIselect size selection (0.6X and 0.8X) on the fragmented cDNA was performed to achieve a fragment range of +/- 200-600 bp. Next, Illumina P5 and P7 adapters were ligated to the size-selected cDNA using the 10X Genomics Adaptor Ligation Mix at 20°C for 15 min. Post ligation cleanup was performed using SPRIselect beads (0.8X), and the ligated cDNA was eluted in 50 µL Buffer EB. Sample index G6 (Dual Index Plate TT Set A, PN-3000431) was added to the cDNA via PCR using the 10x Genomics Library Amplification Mix. Sample indexing PCR was performed with the following conditions: 98°C for 45 sec; 16 cycles of 98°C for 20 sec, 54°C for 30 sec, 72°C for 20 sec; and 72°C for 1 min. post-indexing PCR, libraries underwent a further double-sided SPRIselect size selection (0.6X and 0.8X). Libraries were eluted in 35 µL Buffer EB.

For ATAC library preparation, 500 ng of the SPLONGGET DNA library was used as input to a double-sided SPRIselect size selection (0.6X and 0.8X). A ready-to-sequence ATAC library was eluted in 30 µL Buffer EB. All gene expression and open chromatin libraries were sequenced on an Illumina NovaSeq X Plus.

### Sequencing data preprocessing

After sequencing, pod5 files were basecalled using dorado (v0.7.2.0) with the dna_r10.4.1_e8.2_400bps_sup@v5.0.0 basecalling model. Next, we used two nextflow pipelines, nf-core/scnanoseq^20^ (v1.0.0) and IntGenomicsLabs/scdnalong^21^ (v0.0.1), to perform all quality control steps and processing steps for both the targeted and untargeted transcriptomic and genomic libraries, respectively. Detailed information can be found at https://nf-co.re/scnanoseq/1.0.0/ and https://github.com/IntGenomicsLab/scdnalong. In short, cDNA reads are trimmed using nanofilt^35^ (2.8.0). Following this, barcodes and UMIs were identified using blaze^36^ (2.2.0) providing the 10X_3v3 barcode structure. Next, reads were mapped with Minimap2^37^ (2.28-r1209) to the alt-masked hg38 reference genome, and deduplicated using umitools^38^ (1.1.5). Finally, genes and transcripts were quantified, and novel isoform detection was performed by combining all bam files and running IsoQuant^39^ (3.5.0). For the genomic libraries, barcodes were identified using Flexiplex^40^ (1.0.1) and added to the read name. Following this, reads were mapped using Minimap2^37^ (2.28-r1209) to the alt-masked hg38 reference genome and barcodes were moved from the read name to the BC tag in the aligned bam files using flexiformatter (1.0.2). Finally, we used GATK MarkDuplicates^41^ (3.3.0) to deduplicate reads.

### Single nuclei expression data processing

The generated single nuclei data were processed using a standard transcriptomic pipeline^42^. The raw count matrices per time-point are loaded into memory in AnnData format using Scanpy^43^ (1.10.4) package. AnnData objects per time-point are concatenated prior to any downstream processing. Quality control metrics were calculated for the concatenated AnnData and low-quality cells were filtered out based on 𝑀𝐴𝐷 (mean absolute deviations) automatic thresholding approach, for each quality control metric: 𝑀𝐴𝐷 = 𝑚𝑒𝑑𝑖𝑎𝑛(|𝑋_!_ − 𝑚𝑒𝑑𝑖𝑎𝑛(𝑋)|) with 𝑋_!_being the corresponding metric of an observation. Cells were filtered out if they met any of the following criteria:

1. The *number of genes (log-transformed) with at least 1 count in a cell* is 5 𝑀𝐴𝐷 below or above the 𝑚𝑒𝑑𝑖𝑎𝑛(𝑋).
2. The *sum of counts (log-transformed) for a gene* is 5 𝑀𝐴𝐷 below or above the 𝑚𝑒𝑑𝑖𝑎𝑛(𝑋).
3. The *cumulative percentage of counts for the 20 most expressed genes in a cell* is 5 𝑀𝐴𝐷 below or above the 𝑚𝑒𝑑𝑖𝑎𝑛(𝑋).
4. The *proportion of total counts for a cell which are mitochondrial* is 5 𝑀𝐴𝐷 above the 𝑚𝑒𝑑𝑖𝑎𝑛(𝑋).
5. The *proportion of total counts for a cell which are ribosomal* is 5 𝑀𝐴𝐷 above the 𝑚𝑒𝑑𝑖𝑎𝑛(𝑋).

Since doublet detection with standard tools, such as Scrublet, proved to be ineffective on this data, we performed doublet detection manually by setting a threshold on raw UMI counts. Cells were regarded as doublets if RNA raw UMI counts exceeded 20000 or DNA deduplicated counts exceeded 60000. To account for technical sampling effects during the scRNA-seq experiment, the raw counts were normalized for each cell by the total counts over all genes excluding highly variable genes and then the normalized counts were log-transformed. The top 2,000 highly variable genes were selected using Seurat python implementation through Scanpy and utilized to reduce the dimensionality of the dataset. More specifically, Principal Component Analysis (PCA) with 50 principal components (PCs) was performed followed by Uniform Manifold Approximation and Projection (UMAP) on the 50 computed PCs. Cells were then clustered using the Leiden algorithm^44^ and cell clusters were automatically annotated using CellTypist^45^ (1.6.3). The built-in high-resolution model of immune cell types was utilized and cell types were assigned after over-clustering and majority voting.

### Differential gene expression analysis

From the AnnData object, cellular barcodes corresponding to tumour cells from the four different time points were extracted for downstream differential gene expression analysis. The ‘gene_counts’ matrix files generated by IsoQuant (3.5.0) was then filtered to retain only those barcodes associated with tumour cells at each respective time point. To construct pseudobulk profiles, barcodes for each time point were randomly shuffled and grouped to generate three pseudobulk replicates per time point. Standard DESeq2^46^ v.1.46.0 procedures were then applied to perform differential expression analysis, comparing each time point (treated as one group) against the remaining samples.

### Single-Cell Chromatin accessibility data processing

To obtain reads that represent the standard ATAC part of the data, DNA reads shorter than 1 kb were retained using the SAMtools v1.21 ‘samtools view -e ’length(seq)<1000’ command^47^. The 1 kb threshold was selected based on the distribution of nucleosome footprints. Fragment files were created for each time point from the filtered reads using the SnapATAC2^48^ 2.6.4 function ‘snap.pp.make_fragment_file’. The last column of the resulting fragment files was deleted to assure compatibility with pyCisTopic, using the following command: ‘zcat fragments.tsv.gz | sort -k1,1 -k2,2n | cut -f 1-5 | bgzip > fragments_corrected.tsv.gz’. The fragments were then processed with pyCisTopic^22^ v2.0a0. In short, using the gene expression-based cell type annotation, cell type-specific ATAC pseudobulks were created from the fragment files. These pseudobulks were then used for peak calling with MACS2^49^ v2.2.9.1. The peaks from different cell type pseudobulks were combined and normalized. After quality checking, we retained nuclei with a transcription start site enrichment ≥10 and with ≥1,000 unique fragments. All time points were combined into a single cisTopic object (cell x region count matrix), where regions falling into ENCODE blacklist region list were filtered out^50^. We also removed all potential doublets, that is the cells with DNA counts>60,000 and cDNA counts>20,000. Topic modelling was performed with the default parameters and a model with 25 topics was selected for downstream analysis based on the consensus of ‘Minmo’, ‘log-likelihood’, ‘Arun’ and ‘Cao Juan’ performance metrics. UMAP and t-distributed stochastic neighbor embedding (t-SNE)^51^ dimensionality reduction was performed for visualisation. Leiden clustering was performed with resolution parameter of 0.3. To determine the cell type membership of cells that did not have a gene expression counterpart (i.e. failing gene expression quality control), the cell type was assigned based on the consensus from the k-nearest neighbour cell types with k=5. The accuracy of the method was calculated by masking 20% of annotated cells and comparing their real label with the predicted one. The regions of interest for downstream gene regulatory network analysis were selected by the standard three approaches: topic binarization using the ‘otsu’ method, differentially accessible region analysis, and selection of the top 3,000 regions per topic.

To perform differential accessibility region analysis between time point D0 and Q1, the cells annotated at D0_tumour_cells and Q1_tumour_cells from time points D0 and Q1 were subset from the cisTopic object. Then, the pyCisTopic analysis was rerun for the selected cells, resulting in a cisTopic model with 8 models. Differentially Accessible Region (DAR) analysis was performed with the pyCisTopic built-in Wilcoxon rank-sum test. DARs with log2-fold change > 0.5 and Benjamini-Hochberg adjusted p-value < 0.05 were considered as significant.

To compare short read sequencing annotation with long-read sequencing annotation for time point Q3, pyCisTopic was rerun for all Q3 cells with celltypist annotation generated for this time point only. Model with 6 topics was chosen in this case.

### Gene regulatory network analysis

SCENIC+^22^ (1.0a1) was used to infer gene regulatory networks. First, a transcriptome AnnData object with raw counts was created. Consensus regions from MACS2 peak calling were used to create a custom cisTarget database using ‘create_fasta_with_padded_bg_from_bed.sh’ and ‘create_cistarget_motif_databases.py’ scripts with default parameters. Using transcriptome and chromatin accessibility layers containing AnnData and cisTopic objects respectively, the SCENIC+ Snakemake pipeline was run with default parameters. Only direct TF-to-region-to-gene links were considered and used in downstream analysis and visualisations (i.e., we only consider TFs with experimental evidence for their recognition motifs and not from orthology or motif similarity). For the SCENIC+ run on the D0-Q1 subset of cells, we used the same custom cisTarget database as for the SCENIC+ run on the full dataset. For network visualisation, the gene signature names for the eRegulons and TFs of interest were selected. Genes regulated by TFs of interest were sorted based on triplet rank, and at most the top 50 genes for each eRegulon were selected. Cytoscape (3.10.2) was used for gene regulatory network visualisation.

### Single nuclei copy number profiling

Single-cell copy number profiles of all time points were computed from the SPLONGGET DNA data using ASCAT.sc. Briefly, all genomic reads were counted in 500 kb bins and counts were corrected for GC content using lowess regression. Next, binned read counts were segmented using multipcf using a penalty of 15 on all time points simultaneously. Low quality profiles were filtered out based on hierarchical clustering of the segmented copy number profiles and manual inspection. For RNA-based copy number alteration (CNA) inference, InferCNVpy (0.6.0), a Python reimplementation of inferCNV, was run on the gene-count matrix using default parameters. Normal cells (i.e., non-tumour cells) were used as the reference, serving as the baseline to which expression levels across genomic regions in tumour cells were compared.

### Pseudobulk Whole Genome preparation

For the bulk whole-genome analyses, we generated one matched-normal pseudobulk. Using the gene expression-based annotation we extracted the RNA barcodes of non-tumour cells and matched them to their corresponding DNA counterparts using the 10X Multiome barcode inclusions lists (737k-arc-v1.txt). Reads from normal cells (across all time points) were extracted from the BAM files using SAMtools^47^ (1.21) with the following command: ‘samtools view | grep -F -f normal_tags’. The header was retained separately using: ‘samtools view -H’ and then combined with the filtered reads before converting the final SAM files back into a single BAM file containing the normal reads from all time points. To capture the non-ATAC data, only DNA reads > 1 kb were retained using: ‘samtools view -e ’length(seq) > 1000’’. For the four tumour BAMs, we simply used the original BAM files (including reads from normal cells) and filtered for reads > 1 kb. To phase variants and reads, we first used CLAIR3^52^ (1.0.9) to call single nucleotide polymorphisms (SNPs) at the 1000 Genomes panel of normals loci^53^ on the normal BAM file. Longphase^54^ (1.7.3) was used to phase the resulting VCF and haplotag both the normal and four tumour BAM files.

### Pseudobulk-based variant calling

To obtain bulk copy number profiles for the four tumour time points, we adapted ASCATv (3.1)^23^. First, in the ‘ascat.prepareHTS’ function, the additional flag: additional_allelecounter_flags=”-f 0”’ was set to run enable allelecounter to run on the long-read data. Also, the ‘min_base_qual’ and the ‘minCounts’ parameters were both set to 10. Further the new ‘loci_binsize’ parameter was set to 1200 to reduce correlation between neighbouring SNPs used in copy number segmentation. The segmentation penalty in ‘ascat.aspcf’ was adjusted to 150 to avoid oversegmentation. Finally, ASCAT was run in matched mode for the different time points with the normal BAM as control.

Somatic SVs were called using Severus^55^ (1.1) in multi-sample tumour-only mode. The phased tumour BAM files were used as input along with the phased small variant VCF file. SV calls overlapping Variable Number Tandem Repeat (VNTR) regions in GRCh38 were clustered and normalized using VNTR annotation. Further, a minimum variant support of 6 reads was required Somatic single nucleotide variants (SNVs) were identified using ClairS-TO^56^ (0.4.0) on the original BAM files, without filtering on read length. Variant calling was performed independently for each of the four tumour time points using a minimum coverage threshold of 10. Only SNVs with a PASS filter status were retained for downstream analysis. SNVs detected in two or more time points were included without further filtering. However, for SNVs identified in only a single time point, additional stringency was applied:

1. a quality score (QUAL) > 25
2. a minimum read depth (DP) ≥ 30
3. and an allelic fraction (AF) > 0.15

These criteria ensured high-confidence detection of both recurrent and unique variants.

The final set of SNVs and indels was annotated using GATK Funcotator (4.6.1.0)

### Processing and genotyping of targeted SPLONGGET

To process targeted SPLONGGET data, we used the same pipelines as for the untargeted SPLONGGET (see above). To genotype cells with variants of interest, we ran cellsnp-lite (1.2.3) on the bam files in genotype mode using the whitelist as generated by the pipelines’ output and with ‘minMAF 0.00001’ and ‘minCOUNT 10’. Cells required a total depth ≥ 3 & alt depth ≥ 3 for cDNA and a total depth ≥ 2 and alt depth ≥ 2 for DNA libraries for a variant to be called.

### Calculate log-score gained haplotype based on cn-LOH

To distinguish tumour cells from normal cells and a potential doublet, chromosome 3 was leveraged. Bulk copy number data indicated that tumour cells across all time points exhibited a copy-neutral loss of heterozygosity (cn-LOH) at this chromosome. Under this condition, tumour cells are expected to display only the gained haplotype, whereas normal cells should show both the gained and the lost haplotypes at approximately equal levels.

Pseudobulk profiles were generated separately for CAR-expressing cells within the tumour cluster. From the Q3 tumour pseudobulk, SNP sites were filtered based on their B-allele frequency (BAF) in germline (normal) cells, retaining only sites with BAF values between 0.2 and 0.8 (i.e., heterozygous sites). At each selected site, the allele with the highest read count in the tumour pseudobulk was designated as the gained haplotype (ALT), while the allele with the lowest count was assigned as the lost haplotype. A VCF file was then constructed containing all selected loci with designated ALT and REF alleles according to this classification. Using cellsnp-lite (1.2.0), these variants were genotyped at the single-cell level. For each cell, ALT allele counts were summed across all chr3 loci to generate a TOTAL-ALT score, reflecting the representation of the gained haplotype. The total read depth across these loci was also summed, and the TOTAL-ALT score was subtracted to compute a TOTAL-REF score, representing the lost haplotype. Finally, a log-score for the gained haplotye was calculated for each cell as:

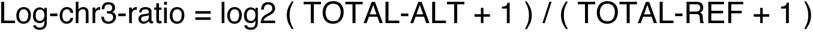

This score quantifies the bias toward the gained haplotype and serves as a discriminative metric for identifying tumour-like cells.

### Doublet detection validation

To validate and detect doublets directly from the raw data, we employed the previous Log-chr3-ratio score for the gained haplotype. This score was plotted against either log10-transformed DNA read counts or RNA UMI counts. Cells with an intermediate Log-chr3-ratio —suggesting a mixture of tumour and normal haplotypes—and simultaneously high RNA or DNA counts were identified as likely doublets. A threshold of 20,000 RNA UMIs or 60,000 DNA reads was applied. Cells meeting these criteria were validated as doublets and excluded from downstream analyses.

### Variant effect prediction

To identify the effects of CD19 intron mutations, we ran Ensembl Variant Effect Predictor (VEP)^27^ v114.2 with the following command: ‘ vep --dir ./vep_data -i cd19.vcf --cache --format vcf --offline –everything’

### De novo genome assembly and identification of CAR construct

To assemble the tisagenlecleucel vector sequence (CTL019; Kymriah), we pulled reads containing part of the lentiviral Long Terminal Repeat from the genome libarary fastqs using flexiplex^40^ (3.5.0) as follows: “*samtools fastq -T ’*’ Q3_genomereads.fastq | flexiplex -x "AGTTAGCCAGAGAGCTCCCAGGCTCAGATCTGGTCT" -d grep -f 3 > CTL019_reads.fastq”*. Flye v2.9.5 was used to run a de novo assembly on these reads via “flye --nano-hq *CTL019_reads.fastq* --genome-size 5k --threads 8 --out-dir $OUTDIR”. The assembled ∼5kb contig corresponding to CTL019 was appended to the alt-masked hg38 reference genome and used for realignment of the cDNA and genome data with Minimap2^37^ (2.28-r1209).

To identify cells with the CAR construct, we selected reads from the bam file (remapped to the reference genome with added CTL019 contig) that mapped to CTL019 contig and extracted and cell barcodes. To identify integration sites, we extracted the primary genomic alignment location and supplementary alignment genomic location from reads mapping to the CTL019 contig.

### Short read library data processing

The short-read sequencing libraries were processed using 10X Genomics CellRanger-ARC (2.0.2) with the prebuilt GRCh38 genome (GRCh38-2020-A-2.0.0) using the following command: ‘cellranger-arc count --reference "${cellranger_arc_reference_dir}" --libraries "${library_file}" --id "${sample_id}" -- localcores 36 --localmem 115 ’.

Downstream analysis of the processed short-read RNA sequencing data was conducted as described in Methods section Single nuclei expression data processing. For processed DNA libraries, the cell type annotations obtained from short read RNA sequencing data output were used to generate cell type-specific pseudobulks from the CellRanger-ARC fragment files. We then followed the same analysis steps as described in Methods section ‘Single Cell Chromatin accessibility data processing’. Model with 15 topics was chosen for downstream pyCisTopic analysis.

### Fusion transcript detection

Fusion transcripts were identified with CTAT-LR Fusion (0.13.0) on the FASTQ files of the four timepoints. The analysis was performed with default settings using the “GRCh38_gencode_v37_CTAT_lib_Mar012021.plug-n-play” CTAT genome library, obtained from the CTAT resources bundle. The top 10 fusions, ranked by the number of supporting reads, and deemed most accurate, are reported here.

## Data availability

The sequencing data has been deposited to EGA under the accession ID XXXX.

## Code availability

Pipelines for preprocessing can be found at https://nf-co.re/scnanoseq/1.0.0/^57^ for transcriptome and https://github.com/IntGenomicsLab/scdnalong^21^. The notebooks and scripts used for the analysis can be found here: https://github.com/IntGenomicsLab/SPLONGGET_analysis.

## Author Contributions

Conceptualization & Methodology: JD. Funding acquisition: JD, JC. Supervision: JD, LH; Data generation: JVV, MA; Resources: HS, MA; Data analysis: JD, LH, AP, RC, ME; Writing – Original Draft: JD, LH, AP, RC, ME, JVV, MA. Writing - Review & Editing: all authors; Visualisation: LH, AP, RC, ME

## Acknowledgements

The compute resources and services used in this work were provided by the VSC (Flemish Supercomputer Center), funded by the Research Foundation - Flanders (FWO) and the Flemish Government. AP is a PhD fellow of the FWO (Grant number: 1159725N). This work was supported by VIB and KU Leuven Internal Funds (C14/22/125 SymBioSys). We would like to acknowledge Koen Theunis, Sarah Geurs, Suresh Kumar Poovathingal, Charalampos (Harry) Anagnostakis, VIB Single Cell Core, VIB Tech Watch, and the Leuven Genomics Core for their assistance and input. We would like to thank Robert A. Forsyth and Laura Beltrame for their help brainstorming method names and logos. We also thank Stein Aerts, Diether Lambrechts, Laurens Lambrechts and Maxime Tarabichi for their critical reading of the manuscript. Figure 1a was created in BioRender (https://BioRender.com/gd9767j)

## Supplementary figures

**Supplementary Fig. 1:**
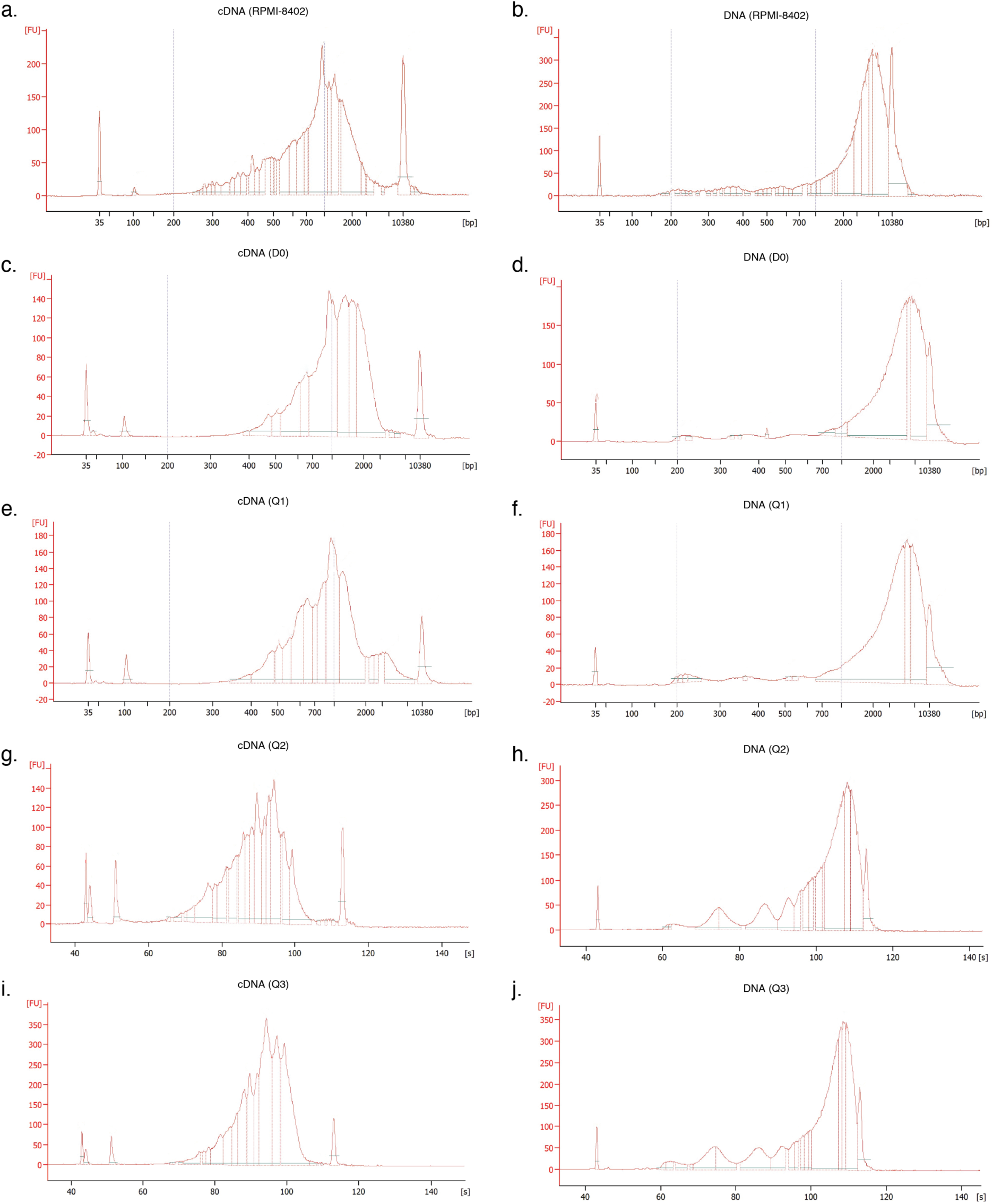
Fragment length distribution of SPLONGGET libraries. **a-j**) Bioanalyzer plot showing fragment lengths in base pairs (bp) of the **a**) RPMI-8402 cDNA, **b**) RPMI-8402 DNA, **c**) D0 cDNA, **d**) D0 DNA, **e**) Q1 cDNA, **f**) Q1 DNA, **g**) Q2 cDNA, **h**) Q2 DNA, **i**) Q3 cDNA, and **j**) Q3 DNA libraries. Note, Bioanalyzer ladders were not captured successfully in **g**-**i**. Therefore, x-axis shows time in seconds (s) instead of fragment size.

**Supplementary Fig. 2:**
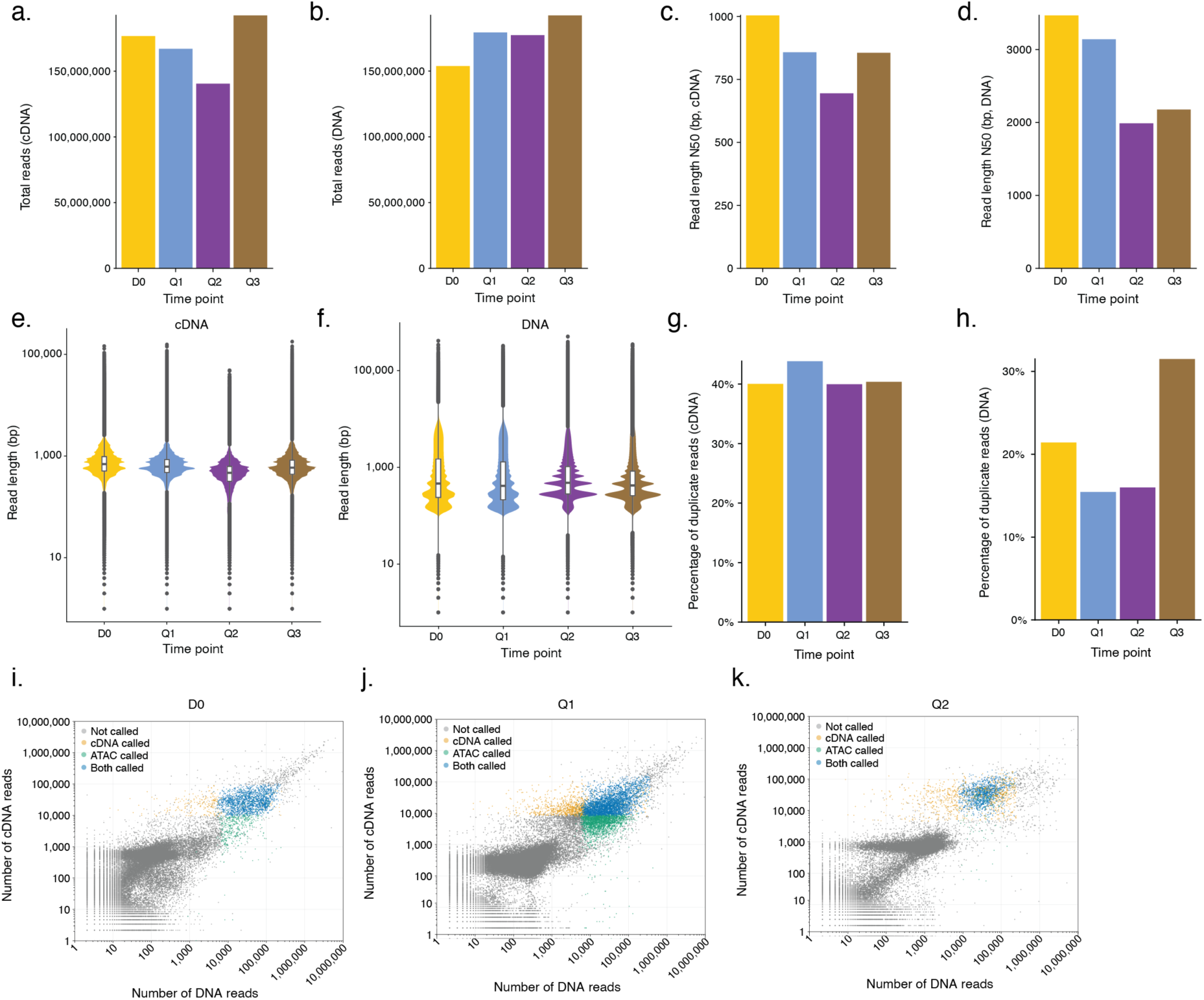
Quality control of sequenced cDNA and DNA libraries. **a-b**) Barplot of total **a**) cDNA and **b**) DNA reads per time point. **c-d**) Barplot of **c**) cDNA and **d**) DNA read length N50s (the length of the shortest read in the set of reads that, when combined, account for 50% of the total bases sequenced) per time point. **e-f**) Violin plot of **e**) cDNA and **f**) DNA read length in base pairs per time point. Boxplot boxes indicate the 25th to the 75th percentile, horizontal bars represent the median, and whiskers extend from –1.5 × IQR to +1.5 × IQR from the closest quartile. IQR, inter-quartile range. **g-h**) Barplot of the percentage of duplicate **g**) cDNA and **h**) DNA reads per time point. **i-k**) Scatter plot of DNA and cDNA counts for **i**) D0, **j**) Q1, and **k**) Q2. Colours depict whether the cell is present as ATAC-only, RNA-only, or is called by both modalities.

**Supplementary Fig. 3:**
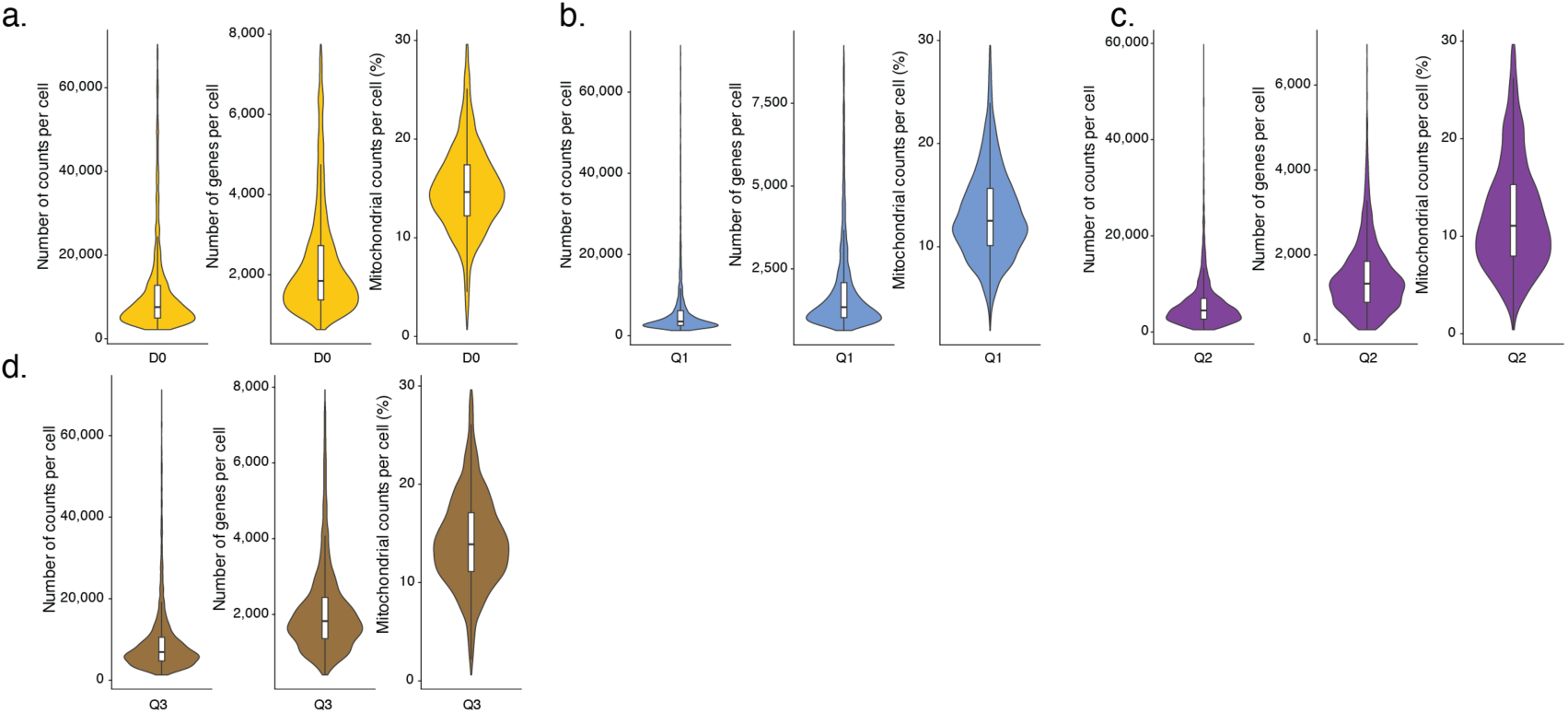
Gene expression data quality control metrics. **a-d**) Violin plots of number of counts per cell, number of genes per cell and percentage of mitochondrial counts per cell for **a**) D0, **b**) Q1, **c**) Q2, and **d**) Q3. Boxplot boxes indicate the 25th to the 75th percentile, horizontal bars represent the median, and whiskers extend from –1.5 × IQR to +1.5 × IQR from the closest quartile. IQR, inter-quartile range.

**Supplementary Fig. 4:**
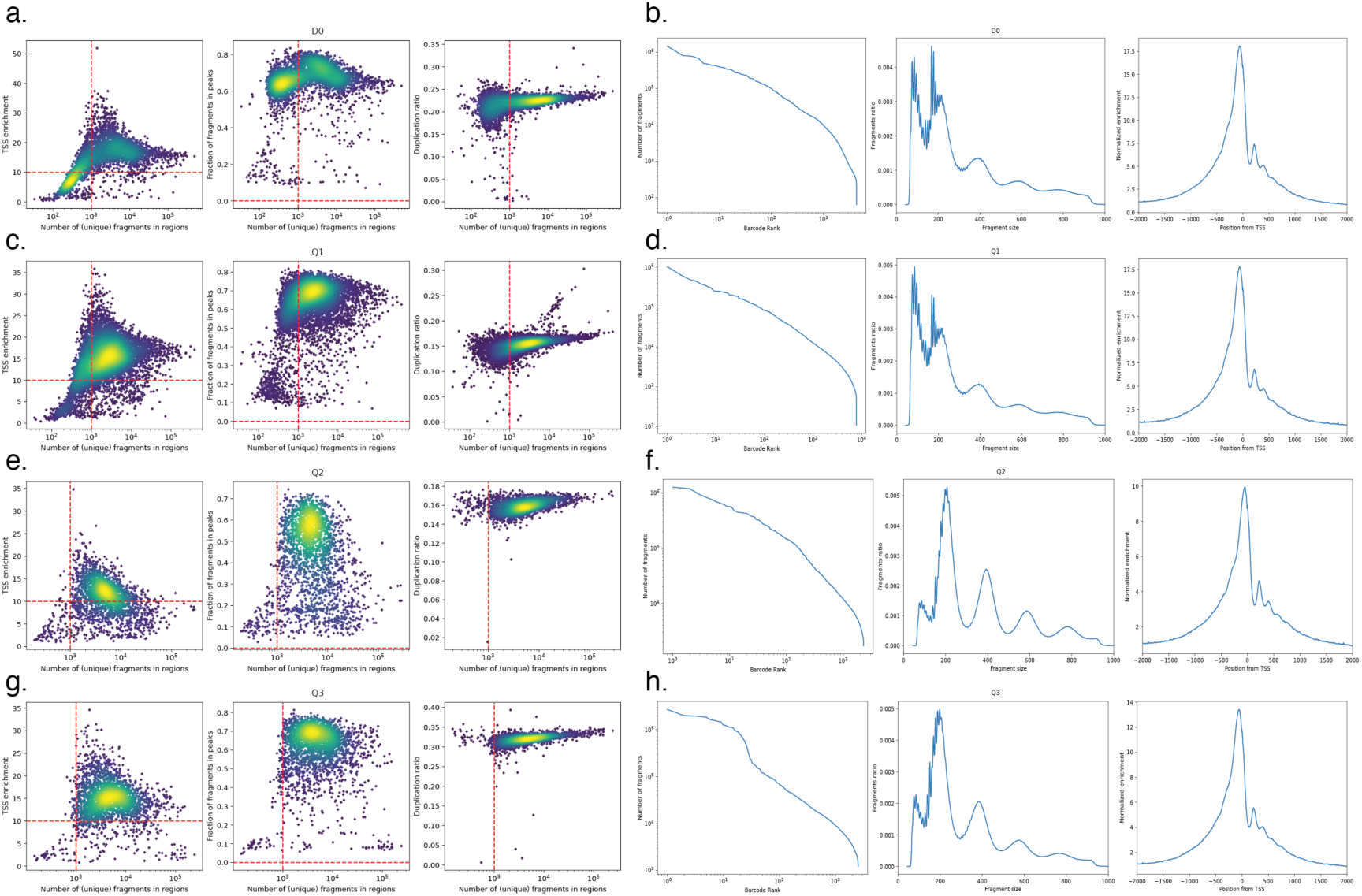
Chromatin accessibility quality control metrics. **a**) Cell barcode statistics for D0. From left to right: number of unique fragments vs TSS enrichment. Number of unique fragments vs fraction of reads in peaks (FRIP). Number of unique fragments vs the duplication ratio. **b**) Sample level statistics for D0. From left to right: Cell barcode rank plot. Fragment size distribution. Normalised TSS enrichment. **c-d**) Same as **a** and **b** but for Q1. **e-f**) Same as **a** and **b** but for Q2. **g-h**) Same as **a** and **b** but for Q3.

**Supplementary Fig. 5:**
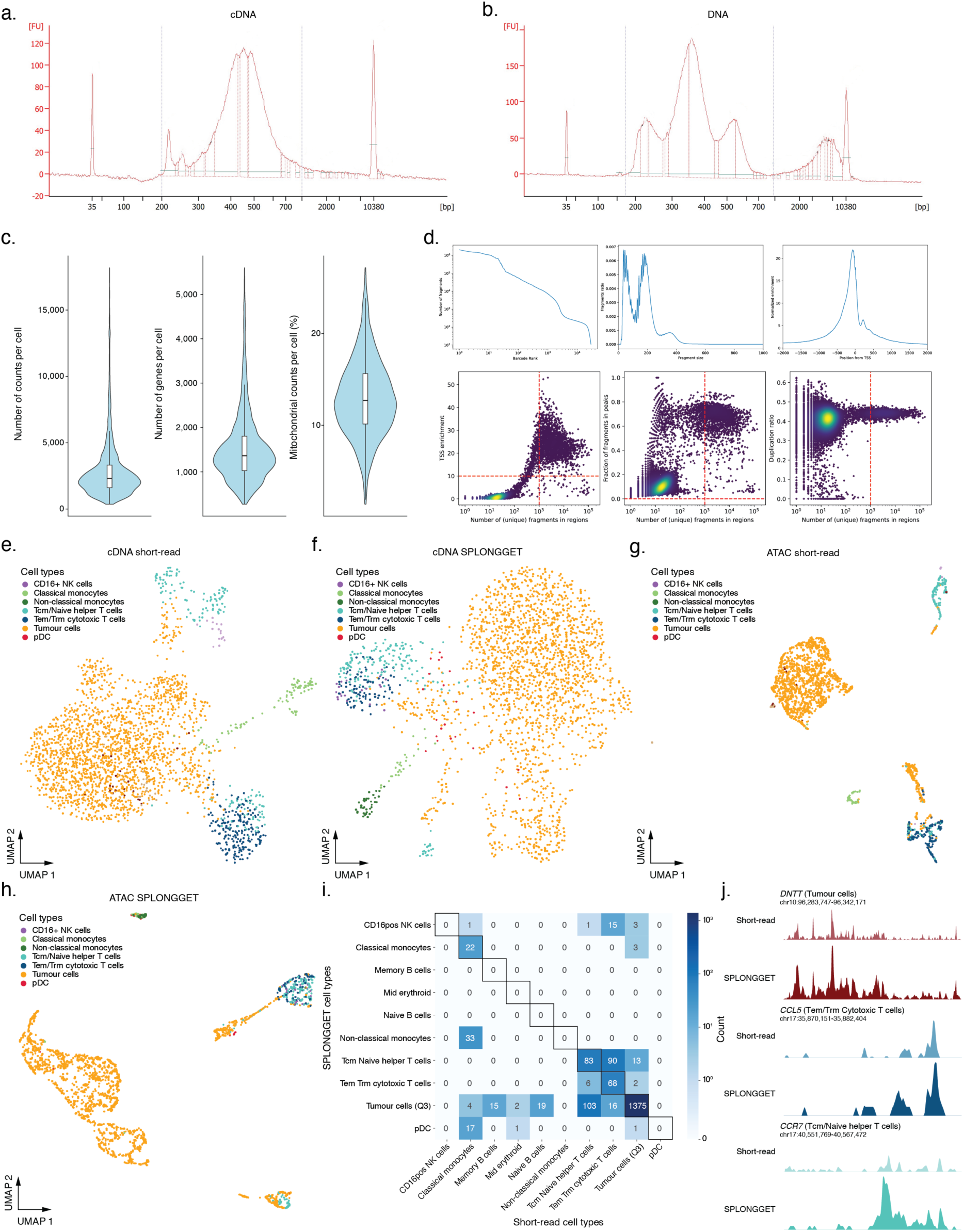
Short-read sequencing of SPLONGGET libraries corroborates gene expression and ATAC read outs. **a-b**) Bioanalyzer plot showing fragment lengths (bp) of the short-read **a**) cDNA and **b**) ATAC SPLONGGET Q3 sublibraries. **c**) Violin plots of QC metrics for short-read Q3 gene expression data. From left to right: read counts, number of genes, and percentage of mitochondrial reads per cell. Boxplot boxes indicate the 25^th^ to the 75^th^ percentile, horizontal bars represent the median, and whiskers extend from –1.5 × IQR to +1.5 × IQR from the closest quartile. IQR, inter-quartile range. **d**) Top: Sample statistics for short-read Q3 ATAC. From left to right: Cell barcode rank plot. Fragment size distribution. Normalised TSS enrichment. Bottom: Cell barcode level statistics for short-read Q3 ATAC. From left to right: number of unique fragments vs TSS enrichment. Number of unique fragments vs fraction of reads in peaks (FRIP). Number of unique fragments vs the duplication ratio. **e-f**) Gene expression UMAP based on **e**) short-read and **f**) SPLONGGET data of Q3. Colours indicate cell types. **g-h**) Chromatin accessibility UMAP based on **g**) short-read and **h**) SPLONGGET data. Colours indicate cell types. **i**) Confusion matrix with cell type annotations of ATAC cells from short-read sequencing vs SPLONGGET. **j**) Normalised BigWig ATAC pseudobulk profiles for regions surrounding marker genes of tumour cells (top two profiles), T_em_/T_rm_ Cytotoxic T cells (middle two profiles), and T_cm_/Naïve helper T cells (bottom two profiles). For each combination, top shows the short-read profile and bottom shows the SPLONGGET profile.

**Supplementary Fig. 6:**
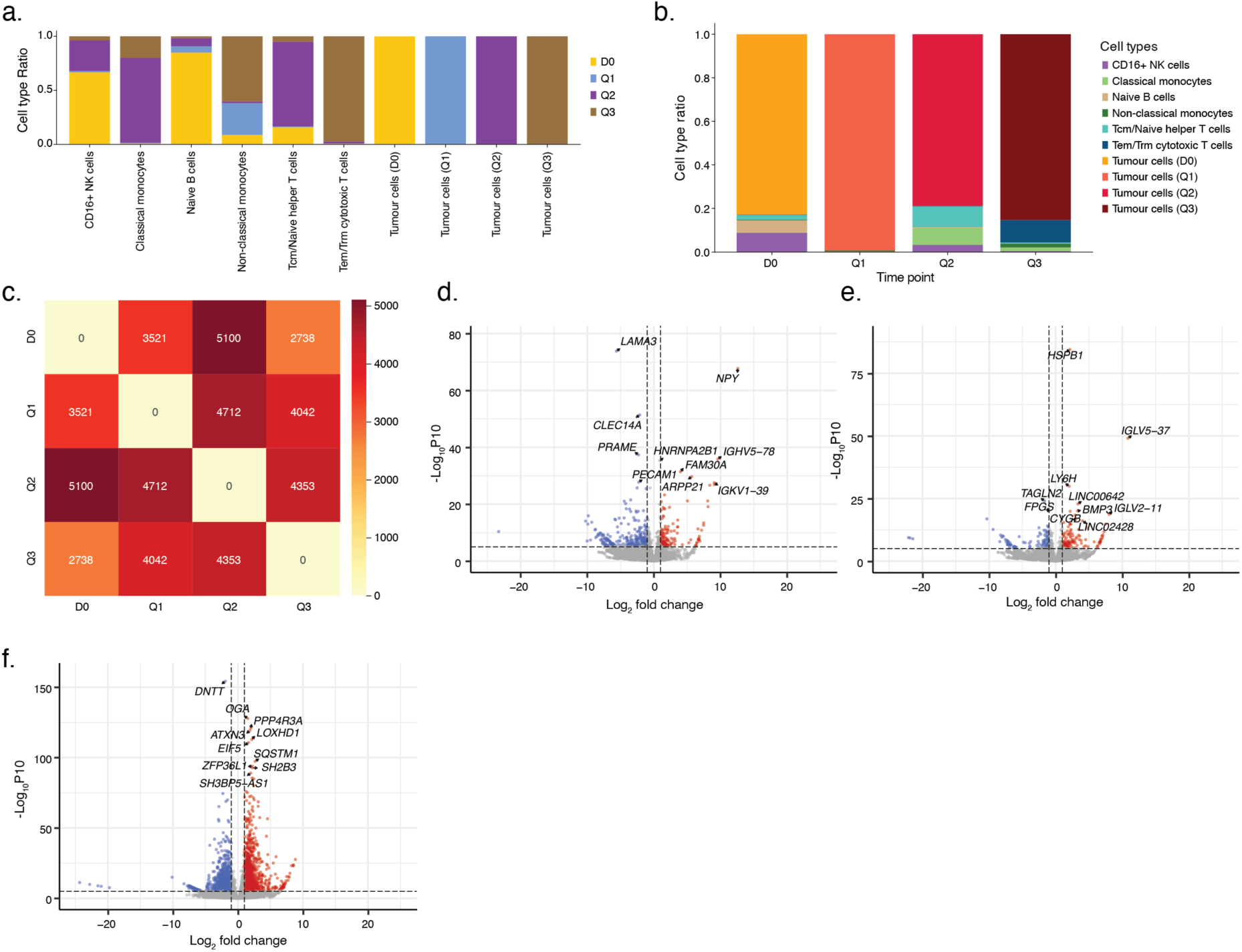
Transcriptome profiling reveals cell type- and time point-specific expression patterns. **a**) Barplots depicting, for each cell type, the fraction of cells derived from each time point. Time points are colour coded. **b**) Barplots depicting, for each time point, the fraction of each cell types. Cell types are colour coded. **c**) Heatmap displaying the number of significantly differentially expressed genes (DEGs) between tumour cell at each pair of time points. **d-f**) Volcano plot showing differentially expressed genes at **d**) D0, **e**), Q1 and **f**) Q2 time points compared to all other time points. The top 10 most significant genes are highlighted.

**Supplementary Fig. 7:**
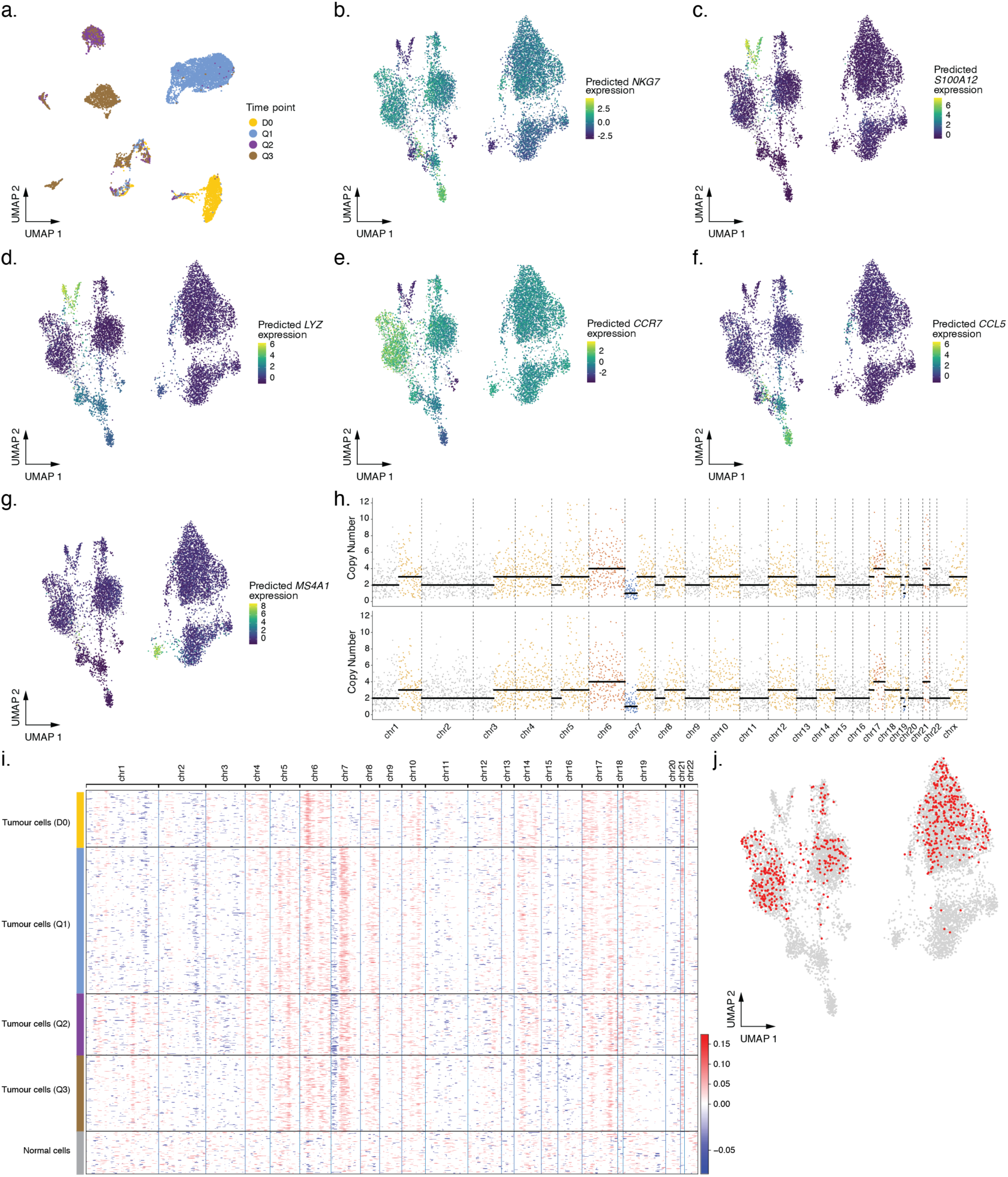
Genomic libraries allow for chromatin accessibility profiling and single-cell copy number profiling. **a**) UMAP of chromatin accessibility data coloured by time point. **b-g**) Predicted gene expression based on chromatin accessibility data for selected marker genes, **b**) *NKG7*, **c**) *S1000A12*, **d**) *LYZ*, **e**) *CCR7*, **f**) *CCL5*, and **g**) *MS4A1.* **h**) Copy number profile of two cells generated by ASCAT.sc. Each coloured dot corresponds to the normalized read count per genomic bin. Black lines indicate copy number segments. Copy number losses are shown in blue, while gains are shown in orange to red. **i**) Heatmap of the copy number landscape inferred by InferCNVpy on the transcriptome data. Each row corresponds to a single cell, and the x-axis represents genomic position. Copy number alterations are shown on a gradient from blue (losses) through white (neutral) to red (gains). Cells are grouped and colour coded by time point. **j**) UMAP showing single cells, with cells highlighted in red indicating detection of isochromosome 7 by single-cell DNA copy number analysis.

**Supplementary Fig. 8:**
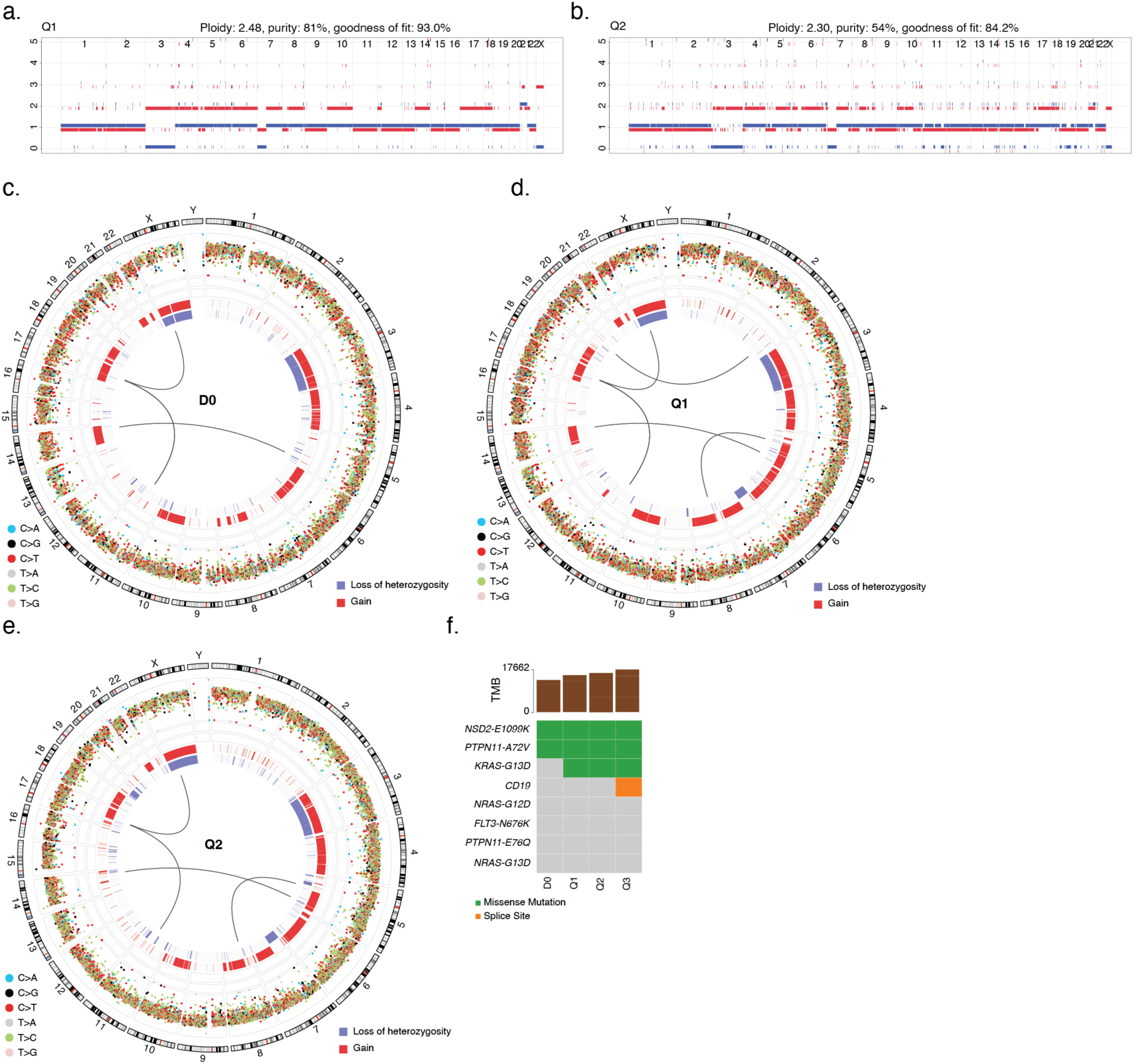
Patient genomic mutational landscape. **a-b**) ASCAT plot showing the allele-specific copy number profile of the **a**) Q1 and **b**) Q2 tumour samples. The red line indicates the major allele copy number, while the blue bars represent the minor allele copy number across the genome. **c-e**) Circos plot of genomic alterations of **c**) D0, **d**) Q1, and **e**) Q2, providing a genome-wide overview of all types of variation. The innermost track displays translocations, with arcs connecting the genomic loci where breakpoints occur. The middle track represents CNAs, where blue bars indicate loss of heterozygosity and red bars indicate copy number gains. The outermost track shows SNVs, colour-coded by substitution class. **f**) Oncoplot displaying mutations of interest across different time points. Each mutation is colour-coded according to the specific variant detected. Tumor mutation burden (TMB) is shown above the plot for each corresponding time point.

**Supplementary Fig. 9:**
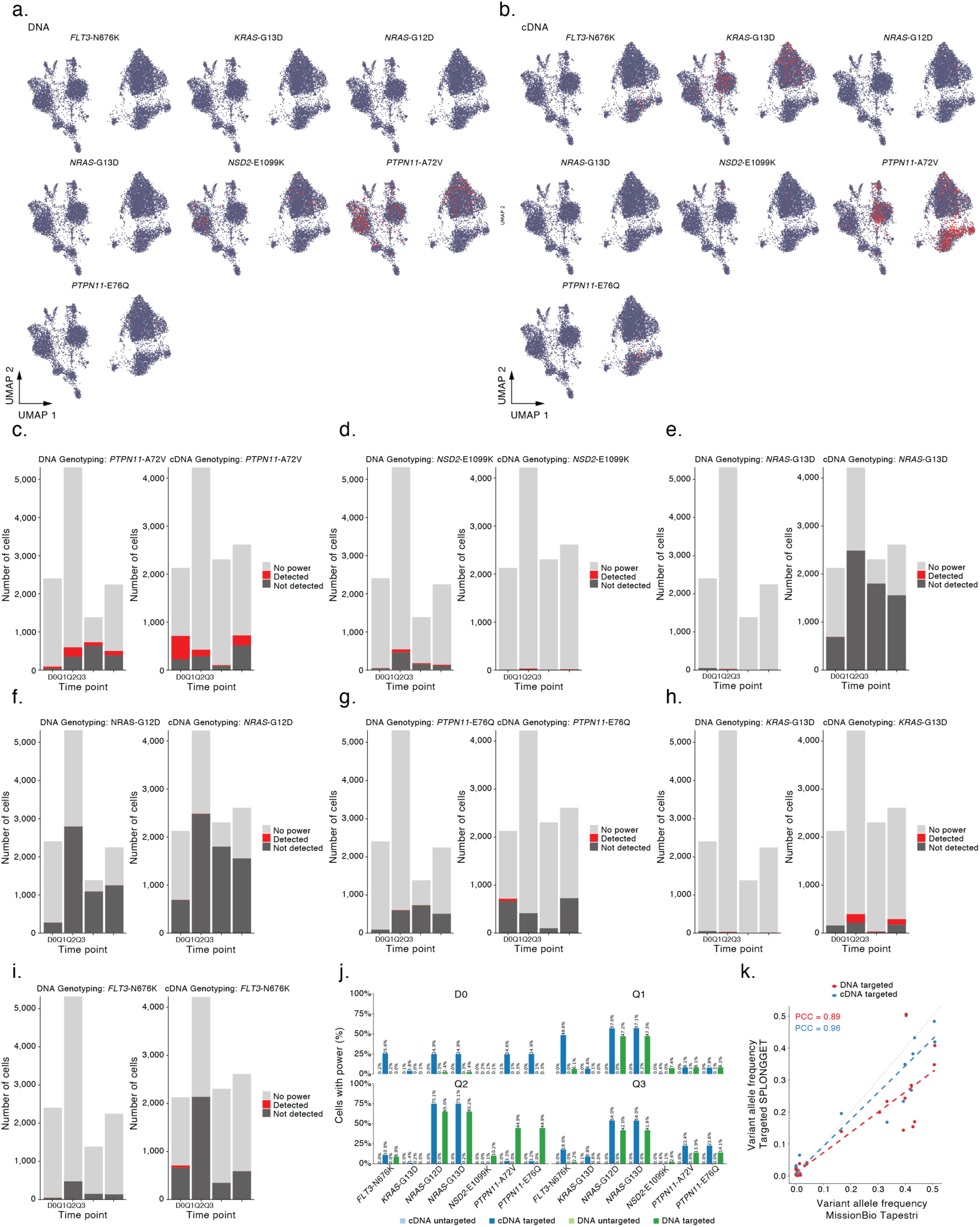
Targeted SPLONGGET enables genotyping variants in single cells. **a-b**) UMAPs showing cells with (red) and without (blue) variant of interest based on **a**) DNA and **b**) cDNA targeted SPLONGGET. **c-i**) Bar plots showing genotyping efficiency of targeted SPLONGGET for **a**) *PTPN11*^A72V^, **b**) *NSD2*^E1099K^, **c**) *NRAS*^G13D^, **d**) *NRAS*^G12D^, **e**) *PTPN11*^E76Q^, **f**) *KRAS*^G13D^, and **g**) *FLT3*^N676K^ across time points, separated by library type: DNA library (left) and cDNA library (right). Each bar represents the number of cells per variant per time point. Bars are colour coded as follows: red for detected variants, black for not detected, and grey for cells lacking coverage to assess the variant. **j**) Bar plots showing the percentage of cells with enough coverage to genotype a targeted variant (total depth ≥ 2 for DNA libraries total depth ≥ 3 for cDNA libraries) for targeted and untargeted DNA and cDNA libraries for the variants of interest. **k**) Dot plot showing the correlation of variant allele frequencies between targeted SPLONGGET (y-axis) and MissionBio Tapestri (x-axis). Red dots: variant allele frequencies from DNA libraries. Blue dots: variant allele frequencies from cDNA libraries. Each dot represents a targeted variant at a specific time point. Dashed red and blue lines: linear regression fits for DNA and cDNA, respectively. PCC: Pearson’s correlation coefficient.

**Supplementary Fig. 10:**
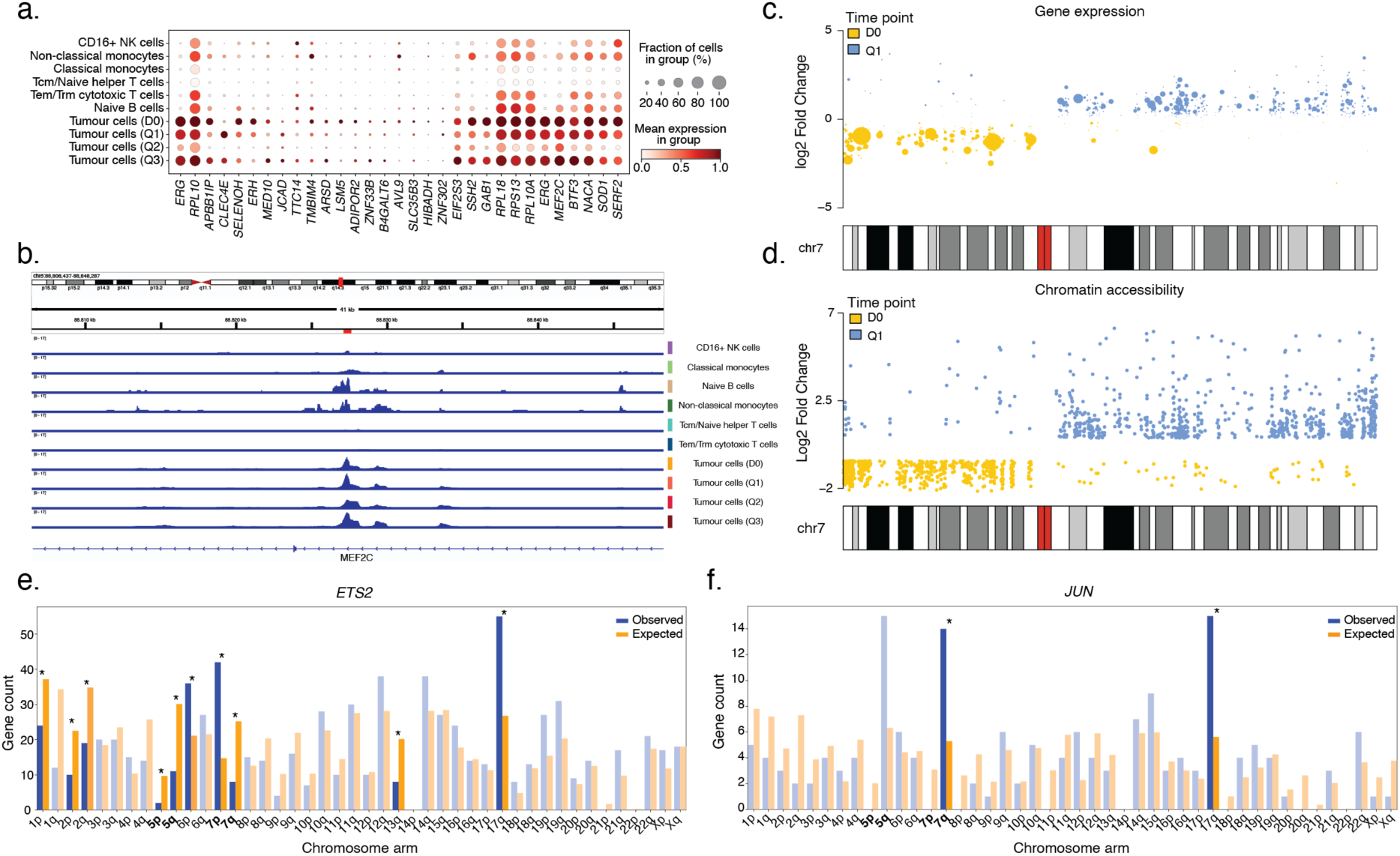
SCENIC+ identifies heterogeneity in relevant gene regulatory networks. **a**) Expression dot plot of *ERG* and its top 30 predicted target genes. Genes were selected based on their triplet rank (TF-region-gene). **b**) IGV screenshot with BigWig ATAC-seq cell type specific pseudobulks spanning *MEF2C*, an *ERG* target gene at chr5:88,806,437-88,848,287. In red, region identified as *ERG* binding region by SCENIC+. **c-d**) Log2-fold change in **c**) gene expression and **d**) chromatin accessibility of genes at chromosome 7 between time point D0 (yellow) and Q1 (blue) tumour cells. **e-f**) Number of expected vs observed number of target genes per chromosome arm for **e**) *ETS2* and **f**) *JUN* eRegulons. Statistical significance is calculated using Chi-square test. * Indicates *P* < 0.05.

**Supplementary Fig. 11:**
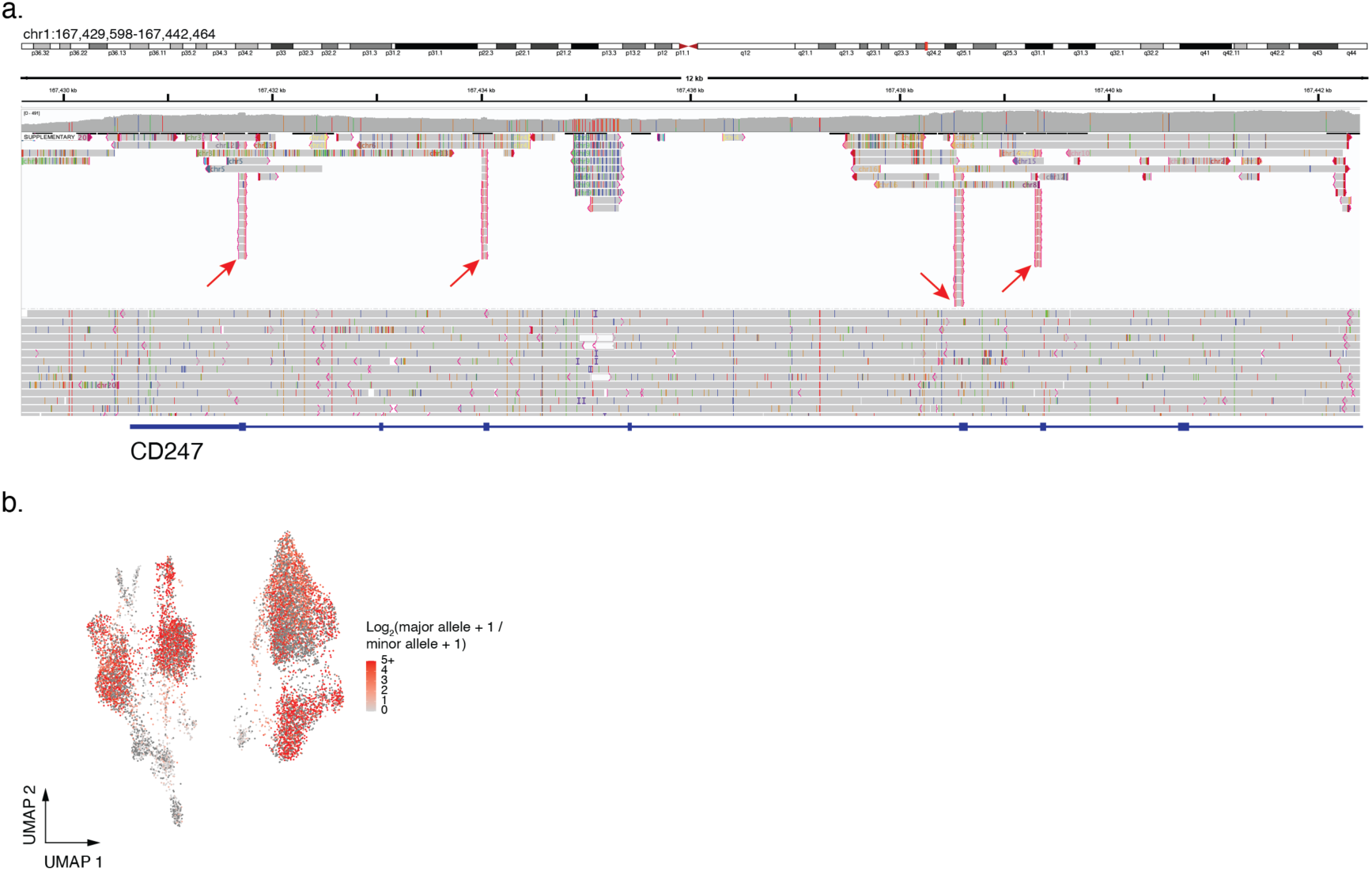
CAR detection in SPLONGGET data. **a**) IGV screenshot of *CD247* with DNA reads from time point Q3. The red arrows highlight the increased coverage across four exons that are part of CAR construct. The reads are clipped since they only map to the exonic part of the gene. **b**) UMAP projection showing the log2-fold read counts for the retained vs lost chr3 haplotype across single cells. A high score identifies tumour cells, while a score around 0 suggests normal (heterozygous diploid) cells. Cells with intermediate scores, shown in light orange, likely represent doublets containing a mix of tumour and non-tumour cells. Dark grey cells indicate cells that did not pass ATAC quality control.

## References

1. Vandereyken, K., Sifrim, A., Thienpont, B. & Voet, T. Methods and applications for single-cell and spatial multi-omics. Nat. Rev. Genet. 1–22 (2023).

2. Buenrostro, J. D. et al. Single-cell chromatin accessibility reveals principles of regulatory variation. Nature 523, 486–490 (2015).

3. Macaulay, I. C. et al. G&T-seq: parallel sequencing of single-cell genomes and transcriptomes. Nat. Methods 12, 519–522 (2015).

4. Zachariadis, V., Cheng, H., Andrews, N. & Enge, M. A Highly Scalable Method for Joint Whole-Genome Sequencing and Gene-Expression Profiling of Single Cells. Mol. Cell 80, 541–553.e5 (2020).

5. Lindenhofer, D. et al. Functional phenotyping of genomic variants using joint multiomic single-cell DNA–RNA sequencing. Nat. Methods 1–10 (2025) doi:10.1038/s41592-025-02805-0.

6. Yu, L. et al. scONE-seq: A single-cell multi-omics method enables simultaneous dissection of phenotype and genotype heterogeneity from frozen tumors. Sci. Adv. 9, eabp8901 (2023).

7. 7. High burden and pervasive positive selection of somatic mutations in normal human skin | Science. https://www.science.org/doi/10.1126/science.aaa6806?url_ver=Z39.88-2003&rfr_id=ori:rid:crossref.org&rfr_dat=cr_pub%20%200pubmed.

8. Izzo, F. et al. Mapping genotypes to chromatin accessibility profiles in single cells. Nature 629, 1149–1157 (2024).

9. Lebrigand, K., Magnone, V., Barbry, P. & Waldmann, R. High throughput error corrected Nanopore single cell transcriptome sequencing. Nat. Commun. 11, 4025 (2020).

10. Inaba, H. & Pui, C.-H. Advances in the Diagnosis and Treatment of Pediatric Acute Lymphoblastic Leukemia. J. Clin. Med. 10, 1926 (2021).

11. Aertgeerts, M. et al. Single-cell DNA and surface protein characterization of high hyperdiploid acute lymphoblastic leukemia at diagnosis and during treatment. HemaSphere 9, e70085 (2025).

12. Brady, S. W. et al. The genomic landscape of pediatric acute lymphoblastic leukemia. Nat. Genet. 54, 1376–1389 (2022).

13. Saleh, K. et al. CAR T-Cells for the Treatment of B-Cell Acute Lymphoblastic Leukemia. J. Clin. Med. 12, 6883 (2023).

14. Laetsch, T. W. et al. Three-Year Update of Tisagenlecleucel in Pediatric and Young Adult Patients With Relapsed/Refractory Acute Lymphoblastic Leukemia in the ELIANA Trial. J. Clin. Oncol. 41, 1664–1669 (2023).

15. Orlando, E. J. et al. Genetic mechanisms of target antigen loss in CAR19 therapy of acute lymphoblastic leukemia. Nat. Med. 24, 1504–1506 (2018).

16. Lamble, A. J. et al. Preinfusion factors impacting relapse immunophenotype following CD19 CAR T cells. Blood Adv. 7, 575–585 (2023).

17. Picelli, S. et al. Tn5 transposase and tagmentation procedures for massively scaled sequencing projects. Genome Res. 24, 2033–2040 (2014).

18. Meyers, S. et al. Monitoring of Leukemia Clones in B-cell Acute Lymphoblastic Leukemia at Diagnosis and During Treatment by Single-cell DNA Amplicon Sequencing. HemaSphere 6, e700 (2022).

19. Ewels, P. A. et al. The nf-core framework for community-curated bioinformatics pipelines. Nat. Biotechnol. 38, 276–278 (2020).

20. Trull, A., Community, N.-C., Worthey, E. A. & Ianov, L. scnanoseq: an nf-core pipeline for Oxford Nanopore single-cell RNA-sequencing. 2025.04.08.647887 Preprint at 10.1101/2025.04.08.647887 (2025).

21. Harbers, L. IntGenomicsLab/scdnalong: 0.0.1. Zenodo 10.5281/zenodo.16099232 (2025).

22. Bravo González-Blas, C., et al. SCENIC+: single-cell multiomic inference of enhancers and gene regulatory networks. Nat. Methods 20, 1355–1367 (2023).

23. Van Loo, P. et al. Allele-specific copy number analysis of tumors. Proc. Natl. Acad. Sci. 107, 16910–16915 (2010).

24. Haas, O. A. & Borkhardt, A. Hyperdiploidy: the longest known, most prevalent, and most enigmatic form of acute lymphoblastic leukemia in children. Leukemia 36, 2769–2783 (2022).

25. Kolmogorov, M., Yuan, J., Lin, Y. & Pevzner, P. A. Assembly of long, error-prone reads using repeat graphs. Nat. Biotechnol. 37, 540–546 (2019).

26. Körber, V. et al. Detecting and quantifying clonal selection in somatic stem cells. Nat. Genet. 1–12 (2025) doi:10.1038/s41588-025-02217-y.

27. McLaren, W. et al. The Ensembl Variant Effect Predictor. Genome Biol. 17, 122 (2016).

28. Kanegane, H. et al. Novel mutations in a Japanese patient with CD19 deficiency. Genes Immun. 8, 663–670 (2007).

29. Venables, J. P. Aberrant and Alternative Splicing in Cancer. Cancer Res. 64, 7647–7654 (2004).

30. Gupta, P., O’Neill, H., Wolvetang, E. J., Chatterjee, A. & Gupta, I. Advances in single-cell long-read sequencing technologies. NAR Genomics Bioinforma. 6, lqae047 (2024).

31. Hu, Y. et al. scNanoATAC-seq: a long-read single-cell ATAC sequencing method to detect chromatin accessibility and genetic variants simultaneously within an individual cell. Cell Res. 33, 83–86 (2023).

32. Grupp, S. A. et al. Chimeric Antigen Receptor–Modified T Cells for Acute Lymphoid Leukemia. N. Engl. J. Med. 368, 1509–1518 (2013).

33. Chen, G. M. et al. Characterization of Leukemic Resistance to CD19-Targeted CAR T-cell Therapy through Deep Genomic Sequencing. Cancer Immunol. Res. 11, 13–19 (2023).

34. Sotillo, E. et al. Convergence of Acquired Mutations and Alternative Splicing of CD19 Enables Resistance to CART-19 Immunotherapy. Cancer Discov. 5, 1282– 1295 (2015).

35. De Coster, W., D’Hert, S., Schultz, D. T., Cruts, M. & Van Broeckhoven, C. NanoPack: visualizing and processing long-read sequencing data. Bioinformatics 34, 2666–2669 (2018).

36. You, Y. et al. Identification of cell barcodes from long-read single-cell RNA-seq with BLAZE. Genome Biol. 24, 66 (2023).

37. Li, H. Minimap2: pairwise alignment for nucleotide sequences. Bioinformatics 34, 3094–3100 (2018).

38. Smith, T., Heger, A. & Sudbery, I. UMI-tools: modeling sequencing errors in Unique Molecular Identifiers to improve quantification accuracy. Genome Res. 27, 491–499 (2017).

39. Prjibelski, A. D. et al. Accurate isoform discovery with IsoQuant using long reads. Nat. Biotechnol. 41, 915–918 (2023).

40. Cheng, O. et al. Flexiplex: a versatile demultiplexer and search tool for omics data. Bioinformatics 40, btae102 (2024).

41. McKenna, A. et al. The Genome Analysis Toolkit: A MapReduce framework for analyzing next-generation DNA sequencing data. Genome Res. 20, 1297–1303 (2010).

42. Heumos, L. et al. Best practices for single-cell analysis across modalities. Nat. Rev. Genet. 24, 550–572 (2023).

43. Wolf, F. A., Angerer, P. & Theis, F. J. SCANPY: large-scale single-cell gene expression data analysis. Genome Biol. 19, 15 (2018).

44. Traag, V. A., Waltman, L. & van Eck, N. J. From Louvain to Leiden: guaranteeing well-connected communities. Sci. Rep. 9, 5233 (2019).

45. Xu, C. et al. Automatic cell-type harmonization and integration across Human Cell Atlas datasets. Cell 186, 5876–5891.e20 (2023).

46. Love, M. I., Huber, W. & Anders, S. Moderated estimation of fold change and dispersion for RNA-seq data with DESeq2. Genome Biol. 15, 550 (2014).

47. Danecek, P. et al. Twelve years of SAMtools and BCFtools. GigaScience 10, giab008 (2021).

48. Zhang, K., Zemke, N. R., Armand, E. J. & Ren, B. A fast, scalable and versatile tool for analysis of single-cell omics data. Nat. Methods 21, 217–227 (2024).

49. Zhang, Y. et al. Model-based Analysis of ChIP-Seq (MACS). Genome Biol. 9, R137 (2008).

50. Amemiya, H. M., Kundaje, A. & Boyle, A. P. The ENCODE Blacklist: Identification of Problematic Regions of the Genome. Sci. Rep. 9, 9354 (2019).

51. Maaten, L. van der & Hinton, G. Visualizing Data using t-SNE. J. Mach. Learn. Res. 9, 2579–2605 (2008).

52. Zheng, Z. et al. Symphonizing pileup and full-alignment for deep learning-based long-read variant calling. Nat. Comput. Sci. 2, 797–803 (2022).

53. Auton, A. et al. A global reference for human genetic variation. Nature 526, 68– 74 (2015).

54. Lin, J.-H., Chen, L.-C., Yu, S.-C. & Huang, Y.-T. LongPhase: an ultra-fast chromosome-scale phasing algorithm for small and large variants. Bioinformatics 38, 1816–1822 (2022).

55. Keskus, A. G. et al. Severus detects somatic structural variation and complex rearrangements in cancer genomes using long-read sequencing. Nat. Biotechnol. 1–11 (2025) doi:10.1038/s41587-025-02618-8.

56. Chen, L. et al. ClairS-TO: A deep-learning method for long-read tumor-only somatic small variant calling. 2025.03.10.642523 Preprint at 10.1101/2025.03.10.642523 (2025).

57. Trull, A., Ianov, L., Brunon, S. & bot, nf-core. nf-core/scnanoseq: nf-core/scnanoseq v1.0.0 - Titanium Toad. Zenodo 10.5281/zenodo.13899280 (2024).

